# In-situ microscopy-assisted meniscus-guided coating for highly sensitive reduced graphene oxide-based nanocomposite biosensor

**DOI:** 10.1101/2024.02.20.580941

**Authors:** Su Yeong Kim, Min Kim, Jeong-Chan Lee, Byeongjoo Jeon, Hanul Kim, Siyoung Q. Choi, Byungkook Oh, Heemin Kang, Hyung-Ryong Kim, Steve Park

## Abstract

Meniscus-guided coating provides great potential for fabricating the nanomaterial-based thin film into high-performance biomedical devices due to the strong relationship between its experimental parameters and the resulting structural properties. However, the complex leverages of various fluid dynamics phenomena hamper optimization of structural properties and device performances. This is due to the absence of in-depth analytical techniques to observe, interpret, and control the solidification process. In this work, we propose an analytical strategy based on the rheological properties of a rGO-based solution using computational fluid dynamics modeling and in situ high-speed microscopy. Through this, we reveal the principles of the solidification mechanism that creates a rGO-based nanocomposite in the form of highly- and evenly-wrinkled thin film and the experimental condition at which this mechanism occurs. The optimized thin film presents high electroconductivity, low chip-to-chip signal variation, and multiplexed electrochemical biosensing performance for three classes of antibodies related to the excessive enrichment of endoplasmic reticulum stress, with detection limits of picomolar levels. This optimizing technique can be universally applied to understanding various solution-based coating systems, and can streamline the production of large-area and high-quality nanocomposite biosensors.

## 1. Introduction

Numerous investigation on nanomaterials has been conducted for their practical applications across diverse biomedical devices, including electronics devices^[1]^, energy storage systems^[2]^, actuators^[3]^, drug delivery^[4]^, and biosensors.^[5]^ To improve the functionality of these devices, nanomaterial-based thin films with extensive structural variability are needed.^[6]^ Moreover, controlling the structural properties through engineering means is particularly important for optimizing the sensitivity and signal accuracy of biosensors. This is because target molecule recognition by capture receptors occurs more prominently on the surface of thin films with improved surface area.

Due to its amorphous carbon structure, reduced graphene oxide (rGO) can form an even broader variety of structures (e.g., wrinkles, interlayer distance, thickness, and porosity) compared to other nanomaterials.^[7]^ For example, wrinkles in rGO-based structures create digitated tunnels that act as individual nanoelectrodes^[6a]^. This not only increases the total geometric surface area, but also improves electroconductivity by electron and mass transfer kinetics^[8]^, and sensitivity by increasing the loading capacity of receptors.^[9]^ Several studies have introduced various coating methods to obtain the wrinkled rGO-based structure, including vacuum filtration^[10]^, hydrothermal coating^[11]^, and drop-casting.^[8b]^ These methods involve applying high temperature and pressure or screening solvent types to adjust solvent evaporation, in order to generate defect-inducing stress to the rGO sheets and thereby form wrinkled structures. Nevertheless, they are limited in maximizing the surface area and biosensing performance of the material, as the structural properties of the resulting nanocomposite are not so diversely alterable by the experimental coating parameters.^[12]^

From this point of view, meniscus-guided coating (MGC) process is acknowledged as an alternative method for crafting rGO-based nanocomposites.^[13]^ Ideally, manipulation of the structural properties should be based on an understanding of the relationship between the coating parameters (e.g., coating speed, substrate temperature, solution concentration) and the fluid dynamics accompanying solidification.^[14]^ However, this raises a challenge due to a variety of intricately jointed fluid dynamics phenomena, such as capillary flow, Marangoni flow, viscous drag, and solvent evaporation components.^[14–15]^ Moreover, the incorporation of polymers to increase the dispersion of rGO molecules (whose one common example is chitosan^[16]^) further complicates the fluid flow behavior. This complexity hinders a complete understanding of the solidification process accompanying coating, which leads to a call for exhaustive trial and error when selecting the combination of experimental parameters to fabricate the rGO-based nanocomposite with the optimal structural properties and biosensing performance. In addition, it degrades the productivity in large-area fabrication and compatibility with industrial manufacturing methods like roll-to-roll processing.^[17]^ To address this, it is necessary to establish an in-depth analytical technique that can investigate the solidification mechanism and the resulting structural properties.

In this work, we propose an approach to interpret the solidification mechanism by combining two analytical technologies: computational fluid dynamics (CFD) modeling and in situ high-speed microscopy, through which we analyzed the rheological properties of the solution near the meniscus and observed the real-time surface configuration at different coating speeds (**Figure 1**). This allows us to understand the principles of controlling the solidification mechanism by changing the coating speed, aimed to optimize the structural properties of the rGO-based thin-film nanocomposite. As a result, we showed that a coating speed of 9 mm s^-1^ resulted in a stable fluid flow behavior and rapid solidification, yielding an ultra-thin nanocomposite with a high number of uniform wrinkles. This thin-film nanocomposite showed enhanced electroconductivity and high electrochemical signal response to target molecules. The comprehension of this structural formation process provides a novel pathway for effectively fabricating a multiplexed and enzymatic method-based electrochemical biosensor for detecting antibodies related to endoplasmic reticulum (ER) stress and relevant chronic diseases.

**Figure 1.**
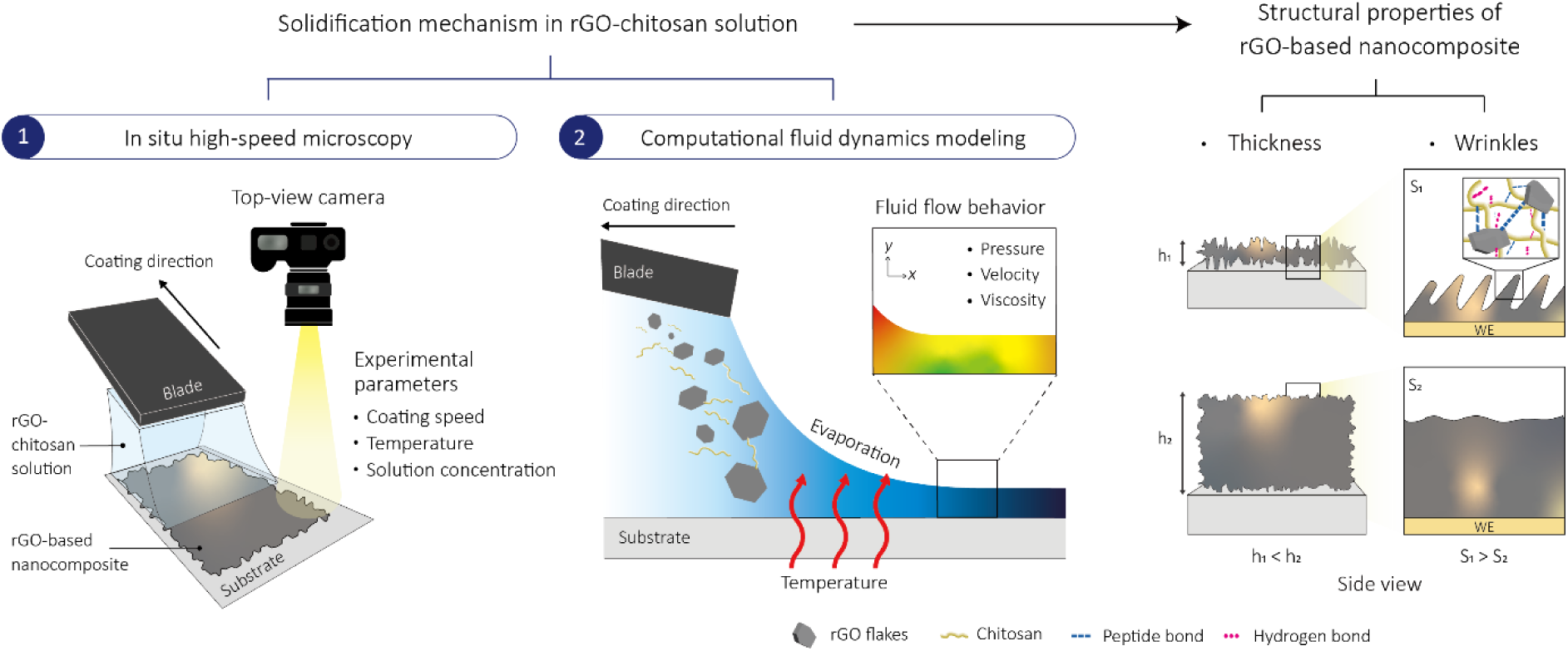
Schematics of investigating the solidification mechanism in solution shearing to optimize the rGO-based nanocomposite through in situ microscopy and computational fluid dynamics (CFD) modeling.

## 2. Results and Discussion

During solution shearing, a chitosan solution in 1% acetic acid dispersed with rGO molecules is dragged across the heated substrate using a blade, continuously depositing the rGO-based nanocomposite near the edge of the meniscus by accelerated solvent evaporation. We first dragged the solution by elevating the coating temperature from 70 °C to 90 °C and by changing the composition and concentration of the rGO-chitosan solution. Comparing the surface coverage of the nanocomposites, we fixed the coating temperature to 90°C and composition of the solution as 5 mg mL^-1^ of rGO for the experiments to follow (**Figure S1** and **S2**, **Supporting information)**. Details were described in **Experimental Section.**

Figure 2a shows the variation of the rGO-based nanocomposite thickness from nanometers to micrometers as the coating speed increases progressively from 0.1 to 100 mm s^-1^. Details in thickness measurement were described in **Figure S3, Supporting information**. Based on the correlation between these two factors, we can divide the results into three different deposition regimes. In the evaporative regime, the meniscus directly affects the rate and extent of coating since the rate of solvent evaporation coincides with the rate of film growth on a similar time scale.^[14a]^ The relationship between the deposited and the incoming solute can be expressed based on mass balance as a power-law (thickness∝coating speed^-1^), which corresponds to an experimental fit with a power of −0.68. In the Landau-Levich (LL) regime, the pressure originates from the back meniscus as the coating speed increases, causing a wet film to be dragged out before solvent evaporation by the increased dominance of the viscous effect.^[14a]^ Thus, the thickness in this regime is influenced by the capillary number (Ca=μν/γ, where μ is solution viscosity, ν is relative velocity, and γ is surface tension of the solution). The fitting result with a power of 0.48 describes the theoretical relationship between the coating speed and thickness in this region (thickness∝coating speed^2/3^). In the mixed regime, which is the transition regime between the evaporative and the LL regime, a thin-film nanocomposite with a minimum thickness of 45.10 nm is achieved at a coating speed of 9 mm s^-1^.

**Figure 2.**
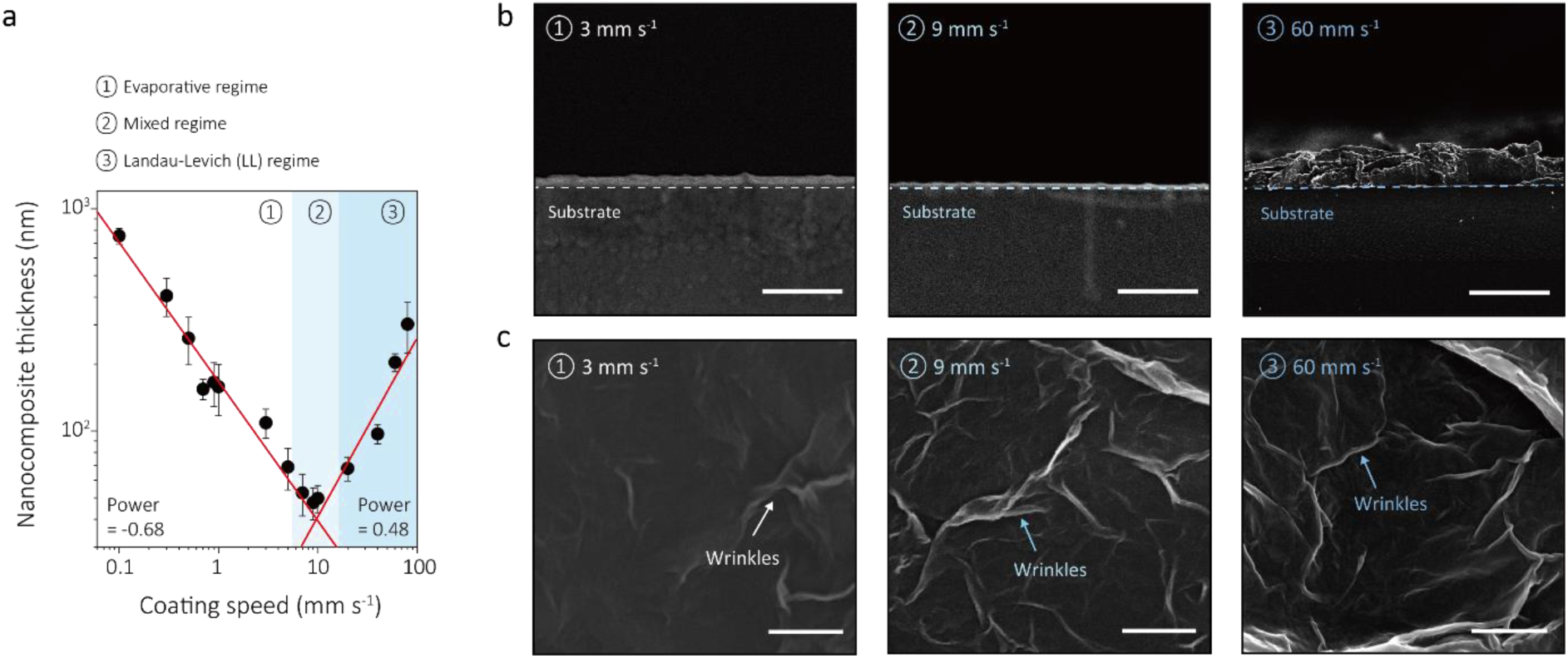
Preparation of the rGO-based nanocomposites with different structural properties. a, Average thickness of rGO-based nanocomposites coated at different coating speeds (n=4, error bars=standard deviation). Red lines represent exponential fits, identifying three different deposition regimes. b, Side and c, Top-view SEM images of rGO-based nanocomposites from three different deposition regimes (3, 9, 60 mm s-1). Scale bars are 500 nm.

Figure 2b shows side-views of SEM images displaying that the thicknesses at three different coating speeds (3, 9, and 60 mm s^-1^) correspond to the results of Figure 2a. Each of the rGO-based nanocomposites belong to the evaporative, mixed, and LL regime, respectively. These nanocomposites will be referred to as by the speed at which they are coated in. Figure 2c shows top-views of these nanocomposites, which in all cases exhibit a high surface coverage but differ in the degree of wrinkling. Here, 9 mm s^-1^ showed a surface packed more densely with wrinkles compared to that of 3 mm s^-1^. This difference was analyzed with Brunaur-Emmett-Teller (BET) surface area measurement, which displayed a 233.73-fold increase in the surface area of 9 mm s^-1^ compared to that of 3 mm s^-1^ (**Figure S4**, **Supporting information**). And despite a 0.92-fold decrease in BET surface area compared to that of 60 mm s^-1^, 9 mm s^-1^ showed a relatively uniform and less agglomerated surface (**Figure S4** and **S5**, **Supporting information**). This difference is apparent from the roughness and the surface profile measurement extracted from AFM analysis (**Figure S5, Supporting information**). From this result, we can confirm that controlling the coating speed leads to changes in both the thickness and wrinkles of the rGO-based nanocomposites. However, the mechanism of this control to optimize the structural properties is ambiguous. To elucidate this, it is important to obtain a deeper understanding of the fluid flow behavior in this coating system. Therefore, we need to interpret the main factors causing changes in the solidification mechanism during solution shearing.

In general, the rheological properties of a solution contribute to the continuous formation of an elongated meniscus, ensuring uniform coating.^[18]^ From this perspective, we would focus on revealing how the rheological properties of the rGO-chitosan solution are manifested in the unique environment of our coating system — shear applied to the solution by the velocity of the blade — and how this alters the fluid flow behavior. To study this, we first conducted CFD modeling dependent on the location (*x*, *y*) values (Figure 3a). In our system, we varied the velocity of the blade (i.e., the coating speed) by setting the substrate to move forward (+*x* direction) at 5, 10, 20, 40, and 60 mm s^-1^ while the blade was stationary (Detailed method is summarized in **Figure S6**-**8** and **Experimental Section, Supporting information**.). For convenience, we will refer to the starting point of the coating as ‘Inside’ or ‘1’ and the finishing point of the coating as ‘Outside’ or ‘2’. The model achieved convergence with a maximum element size coefficient and timespan of 0.005 and 0.03 s, respectively (**Figure S9, Supporting information)**.

**Figure 3.**
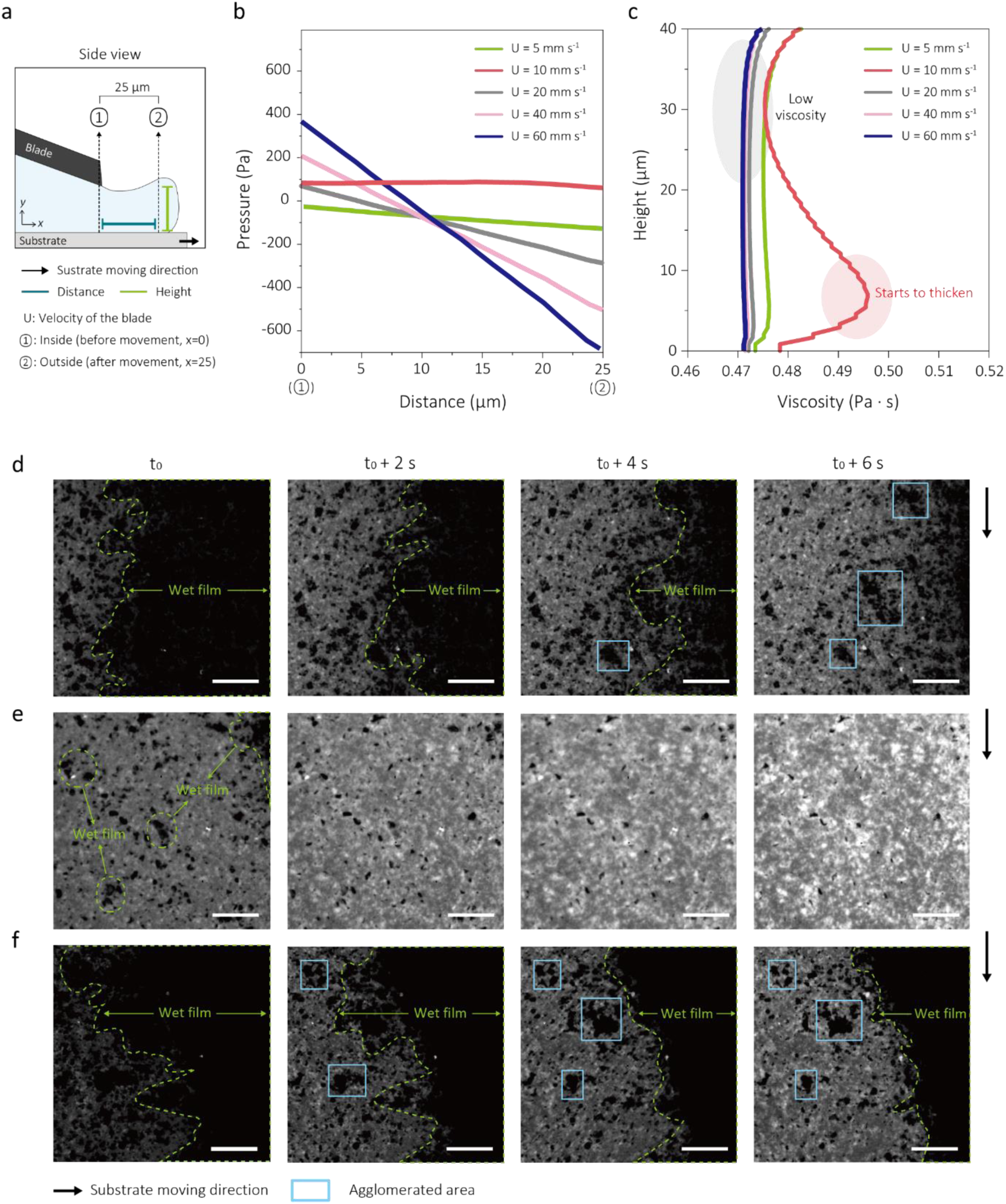
Solidification mechanism investigation by CFD modeling and in situ high-speed microscopy analysis. a, Schematic diagram depicting the parameters used in CFD modeling. b, Pressure profiles of the rGO-chitosan solutions with different velocities of the blade along the +x-direction. The x values range from Inside to Outside. The y values of all data points are 20 μm. c, Viscosity profiles of the rGO-chitosan solutions with different velocities of the blade along the +y-direction. The x-values of the data points are all 25 μm. d-f, Top-view microscopy images of the rGO-chitosan solutions under the coating speed of 3 mm s-1 (d), 9 mm s-1 (e), and 60 mm s-1 (f) as a function of time (Δt=2 s). t0 marks the moment the blade has finished coating the visible area. Black arrows indicate the direction of the substrate movement. Wet film areas are bordered by light green dashed lines. Light blue rectangles indicate the agglomerated areas. Scale bars are 5 μm.

Figure 3b displays the fluid pressure profiles of the rGO-chitosan solutions along the distance between Inside and Outside by changing the velocity of the blade. The fluid pressure is predicted to be constant at 10 mm s^-1^, while the pressures in other velocities rather vigorously decreased along the +*x* direction. This is due to the proportional relationship between the convexity of the fluid velocity along the +*y* direction 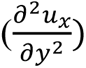 and the gradient of fluid pressure along the +*x* direction 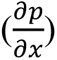 (**Note S1** and **Figure S10, Supporting information)**. This result suggests a constancy in the rheological properties of the solution at 10 mm s^-1^, creating a stable fluid flow behavior and ensuring the elongated meniscus formation. Moreover, it implies that the fluid flow at 10 mm s^-1^ can be maintained without any physical changes. For instance, the fluid thickness at Outside is predicted to remain comparable to that at Inside, whereas the fluid thickness at 60 mm s^-1^ is predicted to increase at Outside compared to the fluid thickness at Inside (**Figure S11, Supporting information)**.

Figure 3c presents viscosity profiles of the fluid located at Outside along the +*y* direction. At 10 mm s^-1^, the viscosity exhibits noticeable thickening at a height of less than 10 µm. Since the rGO-chitosan solution shows a shear thinning behavior (**Figure S8, Supporting information**), this result is attributed to the rapid relief of stress induced by the shear rate, which points to the transfer of kinetic energy between adjacent fluid layers.^[19]^ By contrast, this phenomenon is not observed at the same height for other velocities because the increased fluid thickness resulting from the unstable fluid flow forms additional fluid layers with higher shear rates, and extends the time required for stress relief (**Figure S12, Supporting information**). This difference also indicates that physical changes are minimal at 10 mm s^-1^ and that the fluid flow behavior stabilizes rapidly.

To experimentally verify the fluid flow behavior interpreted by CFD modeling, we conducted a real-time observation of the rGO-based nanocomposite during solution shearing through in situ high-speed microscopy (**Videos S1-3, Supporting information**). Figure 3d-f are the time-dependent top view images of 3, 9, and 60 mm s^-1^ immediately after the solution leaves the tip of the blade (t_0_). The coating speeds of 3 mm s^-1^ and 60 mm s^-1^ were selected from the evaporative and LL regime, respectively, and in the mixed regime, we chose 9 mm s^-1^, which exhibited the smallest thickness (Figure 2a). Comparing the images obtained throughout the 6-second timespan between t_0_ and t_0_ + 6 s, the wet film, observed as the black area, gradually transforms into solidified surfaces, which is shown as the gray area, and the large black dots that remain unchanged are considered as the agglomerated area. At 9 mm s^-1^, the wet film is formed at t_0_, but it dries relatively quickly, with the black droplets mostly transforming into evenly dispersed gray dots (Figure 3e). Since 9 mm s^-1^ falls within the mixed regime, close to the theoretically simulated 10 mm s^-1^ (Figure 2a), this is attributed to constant rheological properties (i.e., low convexity of fluid velocity profile along the +*y* direction, low-pressure gradient along the +*x* direction) and the solidification mechanism where rapid solution thickening occurs. Meanwhile, at 3 mm s^-1^ and 60 mm s^-1^, solidification is considerably slower compared to that of 9 mm s^-1^ (Figure 3d and f). This can be explained by the predicted delay in solution thickening. Also, droplets are severely agglomerated instead of being uniformly dispersed (Figure 3d and f). This droplet-shaped agglomerated area can be explained as the spatial heterogeneity induced by maximized capillary effects due to unstable fluid flow.^[14a]^ For instance, at 3 mm s ^-1^, the stick-and-slip phenomenon can occur and break the meniscus,^[20]^ while at 60 mm s^-1^, the back meniscus acts as an origin of pressure, separating the fluid from the substrate.^[14a]^

These results demonstrate that the stable fluid flow behavior predicted by CFD modeling also operates experimentally under similar experimental parameters, and consequently forms a high and uniformly wrinkled thin-film nanocomposite. That is, the engineering principle between the coating speed, solidification process, and structural properties based on the rheological properties of the solution is successfully established. Based on this principle, the rGO-based nanocomposite exhibits an optimized structure at 9 mm s^-1^.

To confirm the enhanced characteristics of the optimized thin-film nanocomposite, we first conducted cyclic voltammetry at various scan rates in a solution with electroactive species to analyze its electroconductivity. Figure 4a shows that at 9 mm s^-1^, the oxidation and reduction peak currents are attained as high as 80.4% and 92.4%, respectively, of the peak currents of a bare gold electrode. By contrast, 3 mm s^-1^ showed 69.7% and 85.2% of the oxidation and reduction peak currents, and 60 mm s^-1^ only showed 29.2% and 64.3% of the respective peak currents of a bare gold electrode. The correlation between the peak currents and the square root of the scan rate also exhibited a noticeably smaller steepness and weak proportionality for 60 mm s^-1^ compared to those of 3 mm s^-1^ and 9 mm s^-1^ (**Figure S13, Supporting information**). In other words, high structural thickness suppresses the reversibility of oxidation and reduction reactions of electroactive species by increasing the diffusion distance of electrons.^[21]^ 9 mm s^-1^ also showed a distinctively low chip-to-chip variation in electroconductivity with a coefficient of variation (CV) of 5.27% (Figure 4b). This superior consistency in the electrochemical properties of 9 mm s^-1^ is attributed to the highly uniform coating of the nanocomposite without any agglomerated area.

**Figure 4.**
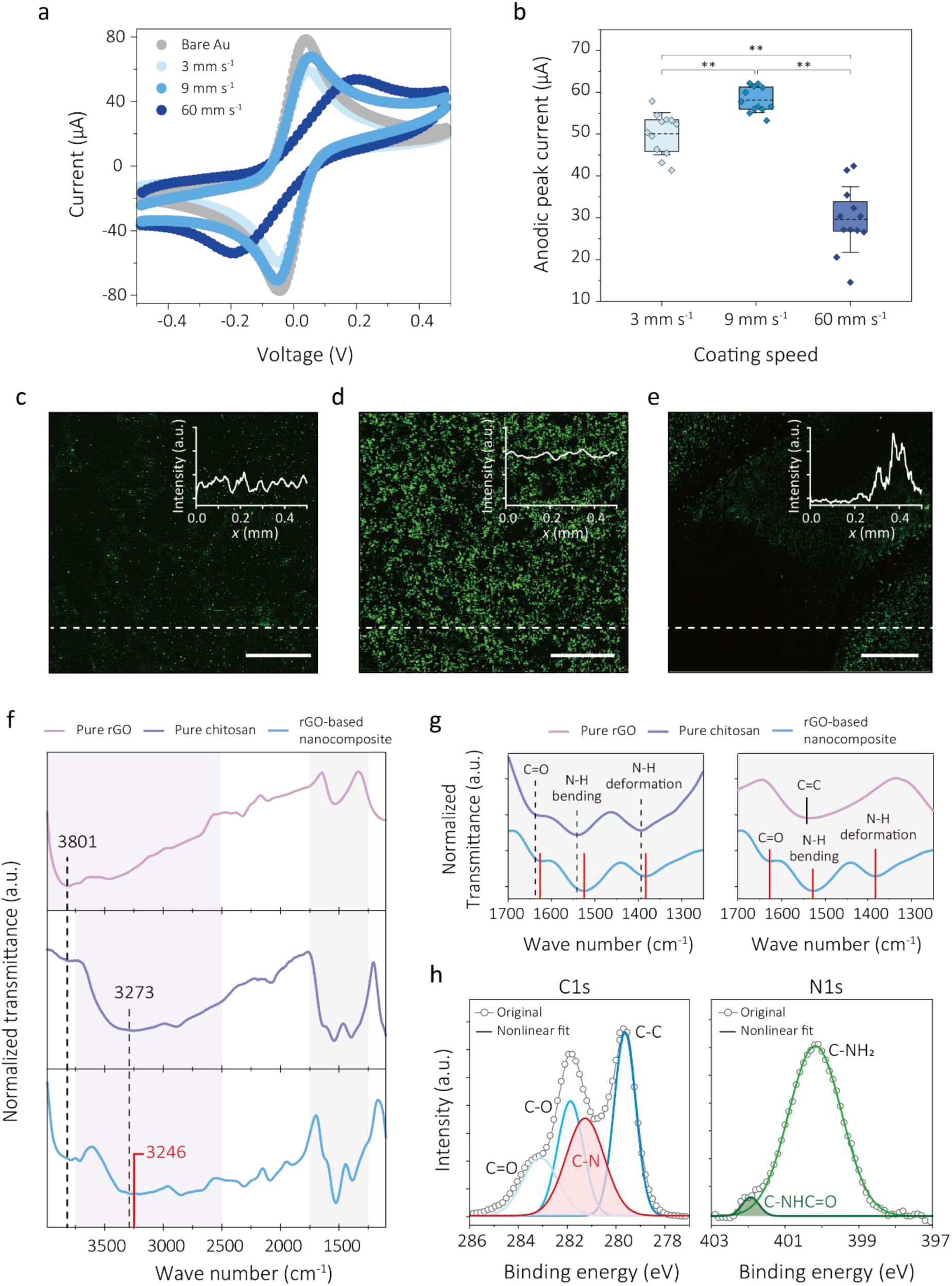
Characterization of the optimized thin-film nanocomposite. a, Cyclic voltammetry profiles showing ferri-/ferrocyanide oxidation and reduction using bare gold electrodes and the optimized nanocomposites (9 mm s-1). Cyclic voltammetry profiles are scanned at 0.2 V s-1, between −0.5 V and 0.5 V. 3 mm s-1 and 60 mm s-1 are used as control samples. b, Peak current distribution of the rGO-based nanocomposites at 3, 9, and 60 mm s-1. (n=12, independent electrode, error bars=standard deviation) Statistical significance was tested (**P < 0.01; one-way ANOVA and tukey post hoc test). c-e, Confocal images of the rGO-based nanocomposite at 3 mm s-1 (c), 9 mm s-1 (d), and 60 mm s-1 (e). All nanocomposites are immobilized with FITC-labeled anti-IgG at excitation wavelength 488 nm. Inset graphs represent the normalized intensity profile along the x-axis (white dashed line). Intensity was measured using Image J. Scale bars are 100 μm. f, Transmittance FT-IR spectra of the optimized nanocomposite compared with rGO and chitosan films. Purple area represents the peaks related to hydrogen bonds and gray area represents the peaks related to peptide bonds. g, Transmittance FT-IR spectra of the chitosan (left) and rGO film (right) each compared to that of the optimized nanocomposite in gray area. h, XPS spectra and their deconvoluted peaks for core C1s (left) and N1s (right) from the optimized nanocomposite. Nonlinear fitting shows R2 values of 0.995 (C1s), and 0.994 (N1s).

To test the functionalization capacity, we then prepared fluorescence analyses by immobilizing FITC-labeled antibodies via EDC/NHS coupling (Figure 4c-e). In this case of 9 mm s^-1^, a confocal image of the nanocomposite displays the green fluorescence of significantly distinctive and consistent clarity with a CV for intensity of 2.43% (Figure 4d). Meanwhile, those of 3 mm s^-1^ and 60 mm s^-1^ show either a low or particularly skewed intensity profiles (Figure 4c**, e** and **Figure S14, Supporting information)**. These notable differences may stem from the high uniformity and degree of wrinkles generated at 9 mm s^-1^. This results in increased loading capacity and leads to an augmented number of receptor binding sites, causing enhancements in sensitivity. The minimal variation in both electroconductivity and fluorescence intensity between chips reduces the variation in electrochemical signals that change by target recognition. Thus, we can anticipate highly sensitive and accurate target detection.

The optimized thin-film nanocomposite also shows successfully formed networks between chitosan and rGO via hydrogen bonds and peptide bonds.^[22]^ First, we observe through Fourier transform infrared spectroscopy (FT-IR) that the O-H stretching peak of the nanocomposite has been red-shifted by 14.6% and 0.82% compared to the corresponding peaks of rGO and chitosan films respectively (Figure 4f). This shifting indicates a weakening of the hydrogen bonds (X-H···X) that were originally present in the pure components, caused by the formation of new additional hydrogen bonds in the form of X-H···Y upon network formation.^[23]^ This is further confirmed by deconvolution of the peaks, showing additional types of hydrogen bonds in the nanocomposite (**Figure S15**, **Supporting information)**. Second, the formation of peptide bonds is exhibited by the red-shift of the involved FT-IR peaks (i.e., C=O, N-H bending, N-H deformation) in the nanocomposite (Figure 4g). This formation is also revealed by X-ray photoelectron spectroscopy (XPS). The C-N and C-NHC=O peak from C1s and N1s spectra indicate successful peptide bonding between the amine groups of chitosan and the carboxylic groups of rGO (Figure 4h and **Figure S16, Supporting information)**. Further spectroscopic data that affirm the successful integration of chitosan within rGO layers including negative and positive time-of-flight secondary ion mass spectrometry (TOF-SIMS), and Raman spectroscopy can be found in **Figure S17** and **S18**, **Supporting information**. These networks enhance the durability of the coating, minimizing the delamination of the coating.^[24]^ This allows the maintenance of the electrochemical signals under wet conditions for 1 week, preserving the shelf-life of the nanocomposite sensor (**Figure S19, Supporting information)**.

As the biosensing paradigm shifts from reactive healthcare to preventive healthcare, the demand for detecting biomolecules that perform a key function in the diagnosis, prognosis, and therapeutic monitoring of chronic diseases has increased.^[25]^ The pathogenesis of chronic diseases involves a common cellular event in which the prolonged unfolded protein response (UPR) fails to mitigate ER stress and activates cell death pathways.^[26]^ In this process, some of the ER stress-related proteins, like glucose-regulating protein 78 (GRP78) and protein disulfide isomerase family A member 6 (PDIA6), are found on the cell membrane of specific cell types or in the serum in their secreted form.^[27]^ These antigenic and serological changes are recognized by the immune system, leading to an autoimmune response, including the generation of circulating autoantibodies (Figure 5a).^[27a]^ Therefore, the autoantibodies in turn can be utilized as crucial and common immunogenic targets to assess the prognosis, treatment, and progression of chronic diseases (e.g., myeloid leukemia, prostate cancer, hepatocellular carcinoma, and ovarian cancer).^[27a, 28]^

**Figure 5.**
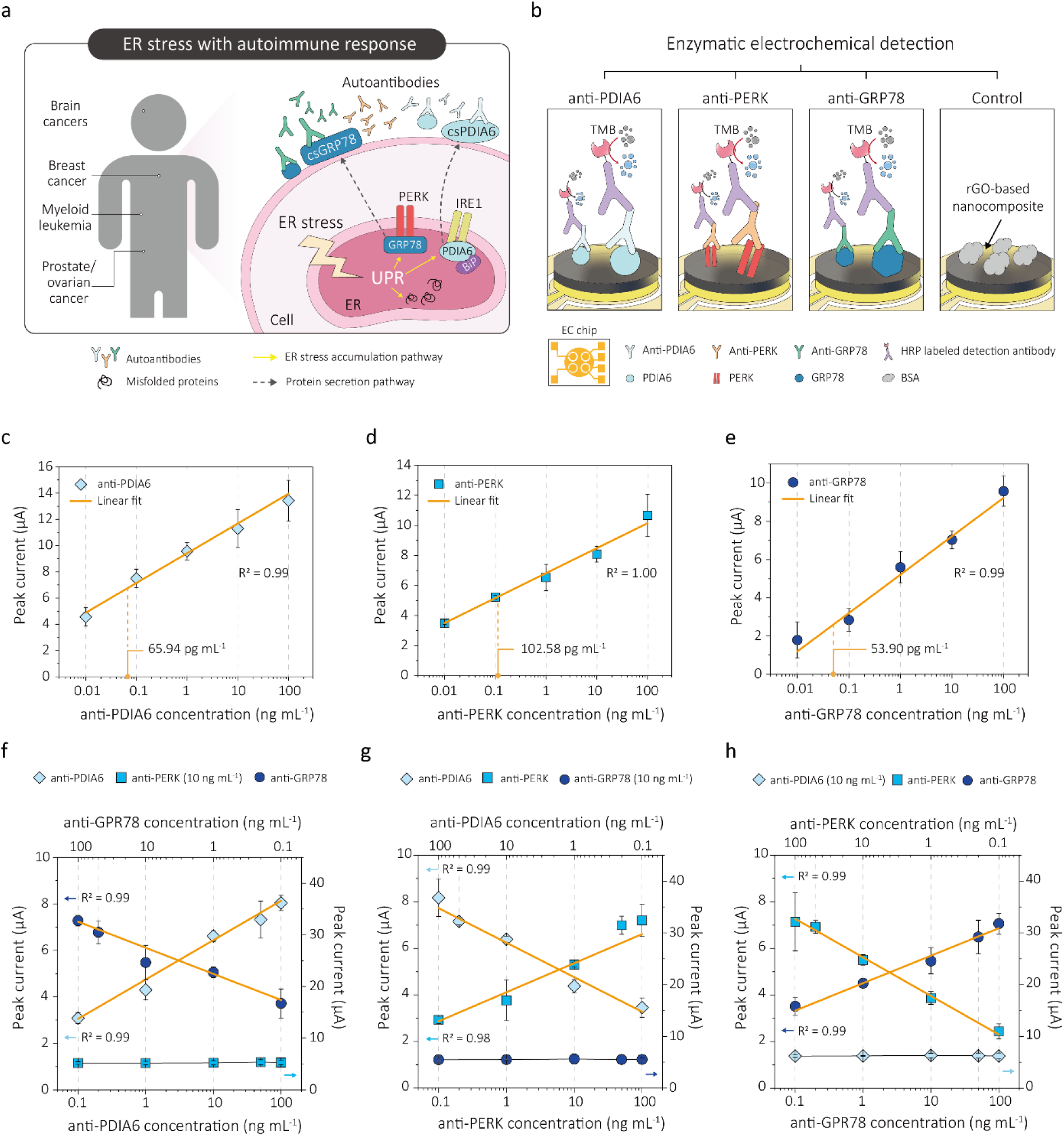
Electrochemical detection of antibodies with the optimized thin-film nanocomposite. **a**, Schematic of endoplasmic reticulum (ER) stress accumulation and related signaling proteins inducing autoimmune response in the human body. **b**, Schematic of the multiplexed electrochemical detection of target antibodies via enzymatic reaction. Target antibodies bind to associated recombinant proteins immobilized to the rGO-based nanocomposite. HRP-labeled detection antibody binding and TMB precipitation follow. Cyclic voltammetry was conducted using four working electrodes, where three are each involved in a reaction with one of the target antibodies, while the one remaining serves as a negative control. **c-e**, Calibration curves for anti-PDIA6, anti-PERK, and anti-GRP78. Peak currents are normalized with peak current values of a blank solution. LOD is defined using three standard deviations (3σ) of the blank solution. (n=4, independent electrodes, error bars=standard deviation) **f**, Calibration curve for the multiplexed detection of anti-PDIA6 concentration (bottom x-axis, left y-axis) and decreasing anti-GRP78 concentration (top x-axis, left y-axis). The concentration of anti-PERK is constant at 10 ng mL^-1^ (right y-axis). **g**, Calibration curve for the multiplexed detection of increasing anti-PERK concentration (bottom x-axis, left y-axis) and decreasing anti-PDIA6 (top x-axis, left y-axis) concentration. The concentration of anti-GRP78 is constant at 10 ng mL^-1^ (right y-axis). **h**, Calibration curve for the multiplexed detection of increasing anti-GRP78 concentration (bottom x-axis, left y-axis) and decreasing anti-PERK concentration (top x-axis, left y-axis). The concentration of anti-PDIA6 is constant at 10 ng mL^-1^ (right y-axis). Peak currents from **f-h** are normalized using peak current values of a blank solution. Yellow lines represent linear fits. Target samples for multiplexed testing were prepared by spiking 10 mM PBS with different concentrations of the three antibodies (n=3, independent electrodes, error bars=standard deviation).

With the optimized rGO-based nanocomposite, we fabricated an electrochemical biosensor and consequently performed an enzymatic-based immunoassay based on the affinity of antibodies and signaling proteins (Figure 5b). First, the target antibody (anti-PDIA6, anti-protein kinase RNA-like endoplasmic reticulum kinase (PERK), and anti-GRP78) binds to its specific capture protein immobilized to the surface. Then, the horseradish peroxidase (HRP)-labeled detection antibody binds to the captured target antibodies, producing surface precipitates when reacted with 3,3’,5,5’-tetramethylbenzidine (TMB) (Figure 5b). The optimization process, where various capture protein and HRP-labeled detection antibody concentrations were tested for this sensor, is further illustrated in **Figure S20** and **S21, Supporting information**.

The performance of this electrochemical sensor was evaluated by classifying the sensors into three types: (1) an anti-PDIA6 sensor, (2) an anti-PERK sensor, and (3) an anti-GRP78 sensor. Subsequently, we immobilized the capture proteins (i.e., recombinant PDIA6, PERK, and GRP78) on each type of sensor to specifically detect their target antibodies. Notably, the calibration curves corresponding to anti-PDIA6, anti-PERK, and anti-GRP78 sensors show a linear increase in peak currents from 100 pg mL^-1^ to 100 ng mL^-1^, and exhibit LOD values as low as 65.94 pg mL^-1^, 102.58 pg mL^-1^, and 53.90 pg mL^-1^ respectively (Figure 5c-e). These detection limits are attributed to increased surface area resulting from the optimization of structural properties (ultra-thin and highly wrinkled structure), and 9 mm s^-1^ indeed exhibited a 2.95- and 2.52-fold increase in target response compared to 3 mm s^-1^ and 60 mm s^-1^, respectively (**Figure S22, Supporting information**).

The high sensitivities of the nanocomposite sensor for anti-PDIA6, anti-PERK, and anti-GRP78, along with the fact that the antibody-target protein pairs do not cross-react with each other, allow us to develop a multiplex electrochemical sensor. This sensor can detect multiple target proteins simultaneously. To do so, three different working electrodes in one sensor were individually immobilized with one of the three different capture proteins, while a fourth electrode was incubated in bovine serum albumin (BSA) solution for control (Figure 5b). To evaluate the sensor’s multiplex performance, we introduced three experimental sets, each consisting of five target samples composed of varying concentrations of anti-PDIA6, anti-PERK, and anti-GRP78, based on their dynamic ranges. The first set consisted of PBS samples spiked with a constant concentration of anti-PERK (10 ng mL^-1^), increasing concentrations of anti-PDIA6 (0.1 – 100 ng mL^-1^), and decreasing concentrations of anti-GRP78 (100 – 0.1 ng mL^-1^). Following the addition of HRP-labeled detection antibodies and TMB precipitation, the peak current signals of anti-PERK exhibited constant values, while signals for anti-PDIA6 increased with increasing concentration, and signals for anti-GRP78 decreased with decreasing concentration, displaying linear calibration curves for anti-PDIA6 and anti-GRP78 respectively (Figure 5f). In the same way, we additionally conducted multiplexed electrochemical detections to produce constant anti-GRP78 signals with decreasing anti-PDIA6 and increasing anti-PERK signals (Figure 5g), and constant anti-PDIA6 signals with increasing anti-GRP78 and decreasing anti-PERK signals (Figure 5h). The exact concentrations and detailed methods can be found in **Table S1, Supporting information** and **Experimental section**. Even though there is a slight signal decrease, the calibration curves reveal consistent linearity similar to what are observed in single-target detection. This finding showcases the applicability of this rGO-based nanocomposite sensor for multiplexed electrochemical detection using biological fluids. Its potentials to uphold specificity for individual targets, and reveal quantitative electrochemical signal changes promise significant advances in the assessment of chronic diseases.

## 3. Conclusion

In this work, we developed a multiplex electrochemical biosensor that can detect three different antibodies (anti-PDIA6, anti-PERK, and anti-GRP78) related to ER stress with the ultra-thin and highly wrinkled rGO-based nanocomposite. Solution shearing is one of the key elements in our work as it finely regulates the structural properties through coating parameters compared to other solution-based processing. Beyond this, we here provide a new analytical technique for this coating method. This technique is assisted by the rheological property analysis through CFD modeling and the real-time observation of surface changes through in situ high speed microscopy. Combining the results of these two analytical tools enables the in-depth investigation of fluid flow behaviors and unexplored solidification processes under different experimental conditions, henceforth allowing us to explain the solidification mechanism and experimentally optimize the structural properties accordingly. This technique will present a simplified path to minimize the extensive amount of simulations required to explain the complex interplay of fluid dynamics components. Furthermore, since this interpretation is focused on elucidating the rheological properties, it can be applied to various types of nanomaterials and coating systems as a reliable guidance for manufacturing large-area and high-quality nanocomposite capable of high-performance biosensors.

Our sensor optimized by this developed technique exhibited enhanced immobilization of capture antigens via carbodiimide reaction, a pronounced reduction in chip-to-chip signal variation, and detection limits of tens to hundreds of pg mL^-1^ with a linear detection range from picomolar to nanomolar level concentrations. Additionally, it showed a capability for the simultaneous detection of three types of antibodies through multiplexed electrochemical sensing. These signal responses in a single, miniaturized chip can be used for understanding cellular health, the progress of chronic diseases, and cancer treatment prognosis. The consolidation of these functions proposes a shift in serological tests for autoantibodies to a point-of-care platform. It will further provide a streamlined solution to the limitations of the current gold standard autoantibody detection methods, such as immunochemistry and western blotting.

## 4. Experimental Section

### Materials

Target antibodies (anti-PDIA6, anti-PERK and anti-GRP78), their corresponding capture antigens (PDIA6, PERK, and GRP78), and horseradish peroxidase (HRP)-labeled goat anti-rabbit IgG as detection antibody were purchased from Abcam (Cambridge, UK). Capture antigens PDIA6 (ab101048), PERK (ab101115), and GRP78 (ab78432) were diluted in 10 mM PBS and used without further purification. Target antibodies for PDIA6 (ab227545), PERK (ab79483), and GRP78 (ab21685), and HRP-labeled detection antibody (ab205718) used for all three target antibodies was diluted in 10 mM PBS. All other chemicals were purchased from Sigma-Aldrich (St. Louis, MO, USA) and were used without further purification.

### Preparation of the rGO-chitosan solution

The rGO-chitosan solution was prepared by dispersing 5 mg mL^-1^ of rGO and 10 mg mL^-1^ of chitosan to 1% (v/v) acetic acid solution in deionized water (DI-water). The solution was sonicated for a total of 30 minutes, with a 30 second on and 10 second off pulse at 20% amplitude (VC 505, Sonics & Materials).

### Solution shearing

The coating blade was fabricated by O_2_ plasma-treated glass substrates at 80 W for 1 minute, followed by (1H,1H,2H,2H-perfluorooctyl) (FTS) treatment through chemical vapor deposition at 70 ℃ for 30 minutes. Substrates were heated at 90 ℃, and an FTS-treated blade was placed at 8 ° angle and 50 µm gap distance to the substrate surface. The rGO-based nanocomposites were coated at coating speeds of 3, 9, and 60 mm s^-1^.

### Rheological properties of the rGO-chitosan solution

The rheological properties of the rGO-chitosan solution were determined with a rheometer (Anton Paar, MCR302). The zero-shear viscosity was measured for various chitosan concentrations (1, 5, 10, and 20 mg mL^-1^) and 5 mg mL^-1^ of rGO suspension in DI-water, using a shear rate of 0.001 s^-1^. Viscosity dependent on shear rate was measured by sweeping the shear rate from 0.01 to 100 s^-1^. Shear stress and viscosity were determined when steady stress was reached. The optimized rGO-chitosan solution (5 mg mL^-1^ of rGO and 10 mg mL^-1^ of chitosan concentration) follows the Carraeau model, where shear rate dependent on viscosity is defined as follows.

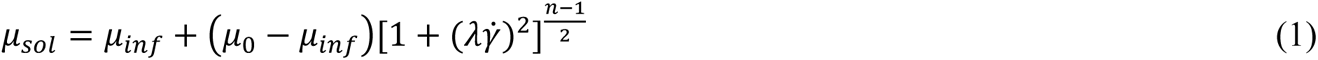

where, μ_*sol*_ is the viscosity, μ_0_ is zero-shear rate viscosity, μ_*inf*_ is infinite shear rate viscosity, *λ* is relaxation time, and n is the power index.

For the control solution, rGO suspension in DI-water was prepared. It follows the Newtonian behavior, where viscosity is constant and independent of 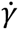.

The surface tension and contact angle of the rGO-chitosan solution were measured through a contact angle analyzer (SEO, Phoenix). Surface tension was analyzed by the pendant drop method, and contact angle was measured at the interface between air, gold electrode, and a single drop (5 uL) of the solution. For comparison, the contact angle of rGO suspension in DI-water was taken.

### Visual and surface characterization of the rGO-based nanocomposites

Top- and side-view of the rGO-based nanocomposites were visualized using scanning electron microscopy (SEM) (Hitachi, S4800). The nanocomposites were coated on O_2_ plasma-treated glass and Si wafers. To measure the thickness, a contact surface profiler (KLA-Tencor, P-15 Long Scan Profiler) was used on the nanocomposites coated on O_2_ plasma-treated Si wafers. Topological images, surface profile and roughness information were obtained through atomic force microscopy (AFM WORKSHOP, PS-2010). For Brunauer-Emmet-Teller (BET) surface area measurement, the rGO-based nanocomposites were prepared into powders first by solution shearing on the glass substrate with a coating speed of 3, 9, and 60 mm s^-1^, followed by annealing at 90 ℃. BET measurement was conducted under N_2_ condition (Micromeritics, ASAP 2420).

### CFD modeling of fluid behavior

To study fluid behavior during solution shearing, a computational fluid dynamics (CFD) modeling was used, utilizing the software COMSOL Multiphysics (COMSOL Inc.) based on the level-set method (LSM) simulation. The initial conditions were set up with the rGO-chitosan solution placed inside the blade and the surrounding areas exposed to air. The simulation proceeded in a time-dependent manner, for a timespan of 0.03 seconds. The Navier slip condition was applied to the rGO-chitosan solution with a coating blade angle of 8 °, and the fluid was assumed to form a fully developed flow with an average velocity (1/2 of the shearing velocity at the outer wall). The following equations were utilized for the system assuming laminar flow (Re<0.006) and incompressible fluid.

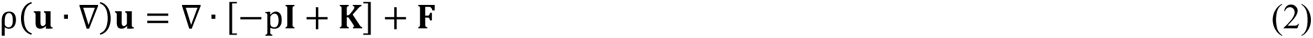

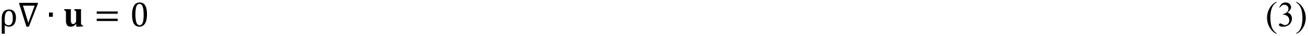

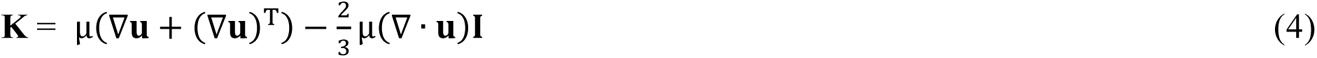

where K is viscous stress tensor, u is fluid velocity, F is external force, ρ is fluid density (assumed as 1000 kg m^-3^), and µ is fluid viscosity.

Equation 1 fitted by Carreau model from the rheological property measurement section was used to account for the shear-thinning behavior of the rGO-chitosan solution: where μ_0_ (zero-shear rate viscosity) was set to 55 Paᆞs, μ_*inf*_ (infinite shear rate viscosity) to 0.47 Paᆞs, *λ* (relaxation time) to 39.8 s, and n (power index) to 0.

An equation representing a two-phase level set was applied to distinguish the boundary between rGO-chitosan solution and air:

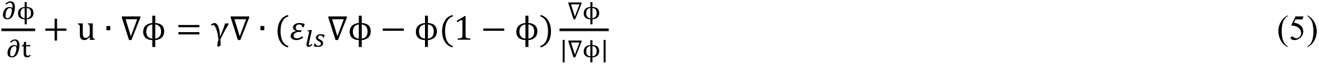

where ϕ is the level set function, γ is reinitialization parameter (1 m s^-1^), and ε0;_*ls*_ is the interface thickness-controlling parameter, or half the maximum element size. The isocontour corresponding to ϕ = 0.5 was designated as the interface between air and the rGO-chitosan solution.

Based on these equations, we measured the velocity profile of the rGO-chitosan solution in the *x*-direction, as well as the pressure and viscosity values near the blade entrance at different velocities (5, 10, 20, 40, and 60 mm s^-1^). The accuracy of the simulation was tested by measuring the convergence with respect to mesh size and simulation time.

### Preparation of the electrochemical (EC) chip

The electrochemical chip consists of 4 working electrodes, 1 reference electrode, and 1 counter electrode. It was produced by depositing metallic source (Cr/Au = 5/50 nm) onto a glass substrate (2.5 cm x 2.0 cm) using an e-beam evaporator (SNTEK, Co., Ltd.). The chips went through a cleaning process involving two steps: a 5 minute sonication in acetone and another 5 minute sonication in isopropanol. Then, the chips were O_2_ plasma-treated at 80 W for 8 minutes (Femtoscience Inc., Korea).

### Electrochemical characterization of the rGO-based nanocomposites

The EC chips coated with the rGO-based nanocomposites at different coating speeds (3, 9, 60 mm s^-1^) were incubated in 50 mM 2-(N-Morpholino) ethanesulfonic acid (MES) buffer containing 400 mM 1-Ethyl-3-(3-dimethylaminopropyl)carbodiimide (EDC) and 200 mM N-hydroxysuccinimide (NHS) for EDC/NHS reaction at room temperature for 30 minutes. Then, the chips were treated with 1 M ethanolamine for 30 minutes in order to quench the unreacted functional groups. Under the electrolyte condition of 5 mM ferri/ferro-cyanide ([Fe(CN)_6_]^3-/4-^) in 1 M KCl, cyclic voltammograms were obtained at scan rate 0.2 V s^-1^ and a scan range of - 0.5 V to 0.5 V (ZIVE SP1, WonATech, Co., Ltd). The oxidation and reduction peak currents at different scan rate were calculated by the software IVMAN 1.5.

### Fluorescence characterization of the rGO-based nanocomposites

The rGO-based nanocomposites produced at different coating speeds (3, 9, 60 mm s^-1^) were functionalized by a solution of 400 mM EDC and 200 mM NHS in a 50 mM MES buffer for 30 minutes. The rGO-based nanocomposites were spotted with FITC-labeled anti-IgG of 1 mg mL^-1^, incubated at 4 ℃ for overnight, and washed thoroughly with 10 mM PBS. Antibody immobilization was visualized through a spinning-disk confocal microscopy system (Andor Dragonfly 200) with an excitation wavelength of 488 nm. Fluorescence intensities were quantified by analysis through Image J.

### Characterization of molecular interactions in the optimized rGO-based nanocomposite

Fourier transform infrared spectroscopy (FT-IR), X-ray photoelectron spectroscopy (XPS), Raman spectroscopy, and both negative and positive time-of-flight secondary ion mass spectroscopy (TOF-SIMS) were conducted to verify formation of an rGO-chitosan network in the optimized rGO-based nanocomposite coated with 9 mm s^-1^. FT-IR spectroscopy was conducted on the networks in their powdered forms, prepared by peeling off the previously coated rGO-based nanocomposite. XPS, Raman spectroscopy, and TOF-SIMS analysis were conducted with the rGO-based nanocomposite coated on O_2_ plasma-treated silicon wafers by solution shearing. For the control sample, drop-casted rGO and chitosan films were used for FT-IR spectroscopy, and drop-casted rGO films were used for XPS and Raman spectroscopy.

### Shelf-life measurement of the optimized rGO-based nanocomposite

The rGO-based nanocomposite coated with 9 mm s^-1^ on the EC chip was incubated with 10 mM PBS at 25 ℃ for following durations: 12 hours, 1 day, and 1 week. Cyclic voltammograms were conducted at scan rate 0.2 V s^-1^ and a scan range of −0.5 V to 0.5 V (ZIVE SP1, WonATech, Co., Ltd) under the electrolyte condition of 5 mM ferri/ferro-cyanide ([Fe(CN)_6_]^3-/4-^) in 1 M KCl. The oxidation and reduction peak currents were calculated by the software IVMAN 1.5.

### Electrochemical detection of anti-PDIA6, anti-PERK, and anti-GRP78

EDC/NHS-treated chips were spotted with 1 µg mL^-1^ of the capture antigens (PDIA6, PERK, and GRP78) on working electrodes (WE1, WE2, WE3) while WE4 was blocked with a solution of 2% BSA in 10 mM PBS. After incubation at 4 °C overnight, the chips were washed with 10 mM PBS, and quenched with 1 M ethanolamine for 30 minutes. They were washed again with 10 mM PBS. The chips were then treated with a blocking buffer of 2% BSA in PBST (0.5 v/v% of Tween 20 in 10 mM PBS) for 1 hour at room temperature. After washing with PBST, the chips were incubated in target antibody solutions or DEPC in the case of blank samples for 30 minutes at room temperature. The chips were washed in PBST and incubated in 50 µg mL^-1^ of HRP-labeled detection antibodies for 30 minutes at room temperature in darkness, once again followed by PBST washing. Lastly, 3,3’,5,5’-tetramethylbenzidine (TMB) was added and incubated for 2 minutes in darkness. Finally, the chips were washed with 10 mM PBS. Using 10 mM PBS as the electrolyte solution, cyclic voltammograms were obtained at a scan rate of 1 V s^-1^ between −0.5 V to 0.5 V (ZIVE SP1, WonATech, Co., Ltd). The oxidation peak currents were calculated by IVMAN 1.5. The limit of detection (LOD) was calculated using the average standard deviation of three blank samples.

### Multiplexed electrochemical detection of anti-PDIA6, anti-PERK, and anti-GRP78

To perform multiplexed detection with one chip, the four working electrodes were each immobilized with 1 µg mL^-1^ of three different capture antigens: PDIA6 on WE1, PERK on WE2, and GRP78 on WE3. 2% BSA solution was spotted on WE4 as control. The working electrodes were incubated in their corresponding solutions at 4 °C overnight. Three different experimental sets consisted of five different target mixtures and were prepared as follows: (1) Increasing concentration of anti-PDIA6 (0.1, 1, 10, 50, 100 ng mL^-1^) spiked in 10 mM PBS with decreasing concentration of anti-GRP78 (100, 50, 10, 1, 0.1 ng mL^-1^) and a constant concentration of anti-PERK (10 ng mL^-1^) (2) Increasing concentration of anti-PERK (0.1, 1, 10, 50, 100 ng mL^-1^) spiked in 10 mM PBS with decreasing concentration of anti-PDIA6 (100, 50, 10, 1, 0.1 ng mL^-1^) and a constant concentration of anti-GRP78 (10 ng mL^-1^), and (3) Increasing concentration of anti-GRP78 (0.1, 1, 10, 50, 100 ng mL^-1^) spiked in 10 mM PBS with decreasing concentration of anti-PERK (100, 50, 10, 1, 0.1 ng mL^-1^) and a constant concentration of anti-PDIA6 (10 ng mL^-1^). These spiked mixtures were added to the chip for 30 minutes. The chips were then washed thoroughly with PBST, and 50 µg mL^-1^ of HRP-labeled detection antibodies were added and incubated for 30 minutes in darkness. The chips were washed with PBST, and they were treated with TMB for 2 minutes in darkness. Finally, the chips were washed thoroughly with 10 mM PBS. Cyclic voltammograms were performed at a scan rate of 1 V s^-1^ with a scan range of −0.5 V to 0.5 V under 10 mM PBS, and oxidation peak currents were calculated by IVMAN 1.5 (ZIVE SP1, WonATech, Co., Ltd).

## Supporting Information

Supporting Information is available from the Wiley Online Library or from the author.

## Acknowledgements

S.Y. Kim and M. Kim equally contributed to this work. This work was supported by the National Research Foundation (NRF) via the Creative Research Initiative Center (Grant No. NRF-2022R1A2C2006076), Institute for Information & Communications Technology Promotion (IITP) grant funded by the Korean government (MSIT, Grant No. 2022-0-00020), and the Ministry of Trade, Industry, and Energy (MOTIE, Grant No. RS-2023-00258591).

---

In-situ microscopy-assisted computational fluid dynamics modeling was employed to explain the effect of the rheological properties of the rGO-based solution on solidification in meniscus-guided coating. Utilizing this analytical technique, the correlation between coating parameters and solidification of reduced graphene oxide was revealed, leading to the development of an optimized uniformly wrinkled ultra-thin nanocomposite with high surface area and electroconductivity as a highly sensitive multiplexed biosensor.

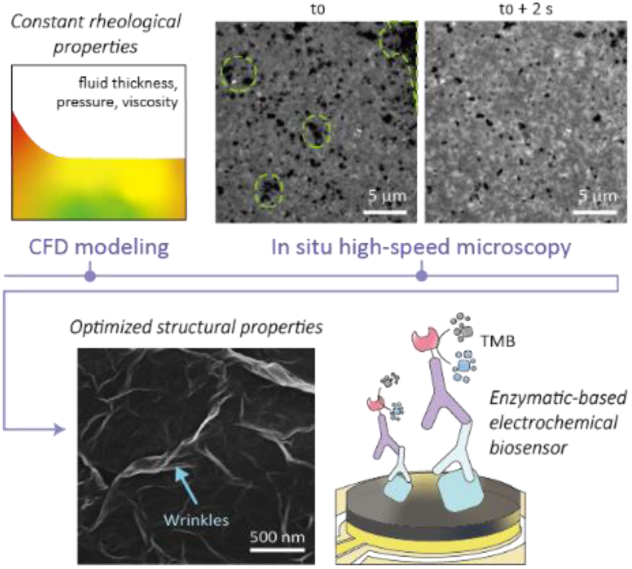

## References

[1] a)T. Lee, J. O. Kim, C. Park, H. Kim, M. Kim, H. Park, I. Kim, J. Ko, K. Pak, S. Q. Choi, I. D. Kim, S. Park, Adv Mater 2022, 34, e2107696; b)J.-C. Lee, J.-O. Kim, H.-J. Lee, B. Shin, S. Park, Chemistry of Materials 2019, 31, 7377; c)D. Viana, S. T. Walston, E. Masvidal-Codina, X. Illa, B. Rodriguez-Meana, J. Del Valle, A. Hayward, A. Dodd, T. Loret, E. Prats-Alfonso, N. de la Oliva, M. Palma, E. Del Corro, M. Del Pilar Bernicola, E. Rodriguez-Lucas, T. Gener, J. M. de la Cruz, M. Torres-Miranda, F. T. Duvan, N. Ria, J. Sperling, S. Marti-Sanchez, M. C. Spadaro, C. Hebert, S. Savage, J. Arbiol, A. Guimera-Brunet, M. V. Puig, B. Yvert, X. Navarro, K. Kostarelos, J. A. Garrido, Nat Nanotechnol 2024.

[2] E. Pomerantseva, F. Bonaccorso, X. Feng, Y. Cui, Y. Gogotsi, Science 2019, 366.

[3] X. Zhu, Y. Hu, G. Wu, W. Chen, N. Bao, ACS Nano 2021, 15, 9273.

[4] a)A. K. Pearce, T. R. Wilks, M. C. Arno, R. K. O’Reilly, Nat Rev Chem 2021, 5, 21; b)Y. J. Kim, B. C. Park, Y. S. Choi, M. J. Ko, Y. K. Kim, Electronic Materials Letters 2019, 15, 471.

[5] a)K. Kim, M. J. Kim, D. W. Kim, S. Y. Kim, S. Park, C. B. Park, Nat Commun 2020, 11, 119; b)S. Y. Kim, J. C. Lee, G. Seo, J. H. Woo, M. Lee, J. Nam, J. Y. Sim, H. R. Kim, E. C. Park, S. Park, Small Sci 2022, 2, 2100111; c)N. M. Naim, H. Abdullah, A. A. Hamid, Electronic Materials Letters 2018, 15, 70.

[6] a)J. Lin, X. Yin, H. Pang, L. Zhang, B. Liao, M. Xin, Y. Su, Y. Liu, L. Wu, K. Jia, B. Zhong, Y. Mai, Y. Dai, Advanced Materials Interfaces 2020, 7; b)J. C. Lee, S. Y. Kim, J. Song, H. Jang, M. Kim, H. Kim, S. Q. Choi, S. Kim, P. Jolly, T. Kang, S. Park, D. E. Ingber, Nat Commun 2024, 15, 711; c)V. Kesler, B. Murmann, H. T. Soh, ACS Nano 2020, 14, 16194.

[7] a)S. Deng, V. Berry, Materials Today 2016, 19, 197; b)J. Kang, F. Li, Z. Xu, X. Chen, M. Sun, Y. Li, X. Yang, L. Guo, JACS Au 2023, 3, 2660.

[8] a)J. N. Chazalviel, P. Allongue, J Am Chem Soc 2011, 133, 762; b)Y. Kang, R. Qiu, M. Jian, P. Wang, Y. Xia, B. Motevalli, W. Zhao, Z. Tian, J. Z. Liu, H. Wang, H. Liu, X. Zhang, Advanced Functional Materials 2020, 30.

[9] M. A. Ali, C. Hu, S. Jahan, B. Yuan, M. S. Saleh, E. Ju, S. J. Gao, R. Panat, Adv Mater 2021, 33, e2006647.

[10] a)W. Zhang, H. Xu, F. Xie, X. Ma, B. Niu, M. Chen, H. Zhang, Y. Zhang, D. Long, Nat Commun 2022, 13, 471; b)G. Zhao, H. Qing, G. Huang, G. M. Genin, T. J. Lu, Z. Luo, F. Xu, X. Zhang, NPG Asia Materials 2018, 10, 982; c)A. Akbari, P. Sheath, S. T. Martin, D. B. Shinde, M. Shaibani, P. C. Banerjee, R. Tkacz, D. Bhattacharyya, M. Majumder, Nat Commun 2016, 7, 10891.

[11] a)P. Zhuang, Z. Guo, S. Wang, Q. Zhang, M. Zhang, L. Fu, H. Min, B. Li, K. Zhang, ACS Omega 2021, 6, 30656; b)T. R. Das, P. K. Sharma, Microchemical Journal 2020, 156.

[12] a)J. C. Lee, H. Seo, M. Lee, D. Kim, H. S. Lee, H. Park, N. Ball, J. Woo, S. Y. Kim, J. Nam, S. Park, Adv Mater 2022, 34, e2105035; b)J. C. Lee, J. H. Woo, H. J. Lee, M. Lee, H. Woo, S. Baek, J. Nam, J. Y. Sim, S. Park, Adv Mater 2022, 34, e2107596.

[13] a)F. Tardani, W. Neri, C. Zakri, H. Kellay, A. Colin, P. Poulin, Langmuir 2018, 34, 2996; b)C. Zhao, P. Zhang, J. Zhou, S. Qi, Y. Yamauchi, R. Shi, R. Fang, Y. Ishida, S. Wang, A. P. Tomsia, M. Liu, L. Jiang, Nature 2020, 580, 210.

[14] a)X. Gu, L. Shaw, K. Gu, M. F. Toney, Z. Bao, Nat Commun 2018, 9, 534; b)G. Giri, E. Verploegen, S. C. B. Mannsfeld, S. Atahan-Evrenk, D. H. Kim, S. Y. Lee, H. A. Becerril, A. Aspuru-Guzik, M. F. Toney, Z. Bao, Nature 2011, 480, 504.

[15] R. Janneck, F. Vercesi, P. Heremans, J. Genoe, C. Rolin, Adv Mater 2016, 28, 8007.

[16] H. Zhang, H. Qu, J. Cui, L. Duan, RSC Adv 2022, 12, 25844.

[17] H. Sun, Q. Wang, J. Qian, Y. Yin, Y. Shi, Y. Li, Semiconductor Science and Technology 2015, 30.

[18] a)G. H. Lee, Y. R. Lee, H. Kim, D. A. Kwon, H. Kim, C. Yang, S. Q. Choi, S. Park, J. W. Jeong, S. Park, Nat Commun 2022, 13, 2643; b)J. C. Lee, M. Lee, H. J. Lee, K. Ahn, J. Nam, S. Park, Adv Mater 2020, 32, e2004864.

[19] F. Bossler, L. Weyrauch, R. Schmidt, E. Koos, Colloids Surf A Physicochem Eng Asp 2017, 518, 85.

[20] W. Han, Z. Lin, Angew Chem Int Ed Engl 2012, 51, 1534.

[21] N. Elgrishi, K. J. Rountree, B. D. McCarthy, E. S. Rountree, T. T. Eisenhart, J. L. Dempsey, Journal of Chemical Education 2017, 95, 197.

[22] S. Kim, M. Cho, Y. Lee, Advanced Functional Materials 2020, 30.

[23] J. Tian, D. Fu, Y. Liu, Y. Guan, S. Miao, Y. Xue, K. Chen, S. Huang, Y. Zhang, L. Xue, T. Chong, P. Yang, Nat Commun 2023, 14, 2816.

[24] a)Y. Jin, W. Lee, Langmuir 2019, 35, 5427; b)Z. Li, S. Gadipelli, H. Li, C. A. Howard, D. J. L. Brett, P. R. Shearing, Z. Guo, I. P. Parkin, F. Li, Nature Energy 2020, 5, 160.

[25] P. Li, G. H. Lee, S. Y. Kim, S. Y. Kwon, H. R. Kim, S. Park, ACS Nano 2021, 15, 1960.

[26] D. R. Peter Walter, Science 2011, 334, 1081.

[27] a)I. Hernandez, M. Cohen, Cancer Lett 2022, 524, 1; b)C. Fonseca, R. Soiffer, V. Ho, M. Vanneman, M. Jinushi, J. Ritz, D. Neuberg, R. Stone, D. DeAngelo, G. Dranoff, Blood 2009, 113, 1681; c)N. S. A. Rahman, S. Zahari, S. E. Syafruddin, M. Firdaus-Raih, T. Y. Low, M. A. Mohtar, Cell Biosci 2022, 12, 129.

[28] a)H.-Y. Kim, J.-H. Lee, M. J. Kim, S. C. Park, M. Choi, W. Lee, K. B. Ku, B. T. Kim, E. C. Park, H. G. Kim, Biosensors and bioelectronics 2021, 175, 112868; b)A. Raiter, A. Vilkin, R. Gingold, Z. Levi, M. Halpern, Y. Niv, B. Hardy, Int J Biol Markers 2014, 29, e431; c)S.-x. H. Xia Ying, Chen-chen He, Cong-ya Zhou, Yi-ping Dong, Meng-jiao Cai, Xin Sui, Cheng-xian Ma, Xiao Sun, Yuan-yuan Zhang, Wen-li Gou, Clifford Mason, Qing Zhu, Oncotarget 2017, 8, 24828.

