## Supporting information for "In-situ microscopy-assisted meniscus-guided coating for highly sensitive reduced graphene oxide-based nanocomposite biosensor"

**Supplementary Note S1.**

When a fluid shows laminar flow under steady state, the relationship between the fluid's speed and pressure can be expressed through the following incompressible Navier-Stokes equation:

$$\nabla \mathrm{p} = \mu \nabla^2 u \tag{1}$$

where p is pressure, and $u$ is the speed of the fluid.

Assuming that the substrate and blade are parallel, we can approximate that

$$\frac{\partial^2 u_x}{\partial y^2} >> \frac{\partial^2 u_x}{\partial x^2} \approx 0 \ (Lubrication\ approximation) \tag{2}$$

With this assumption, the *x* component from Equation 1 can be expressed as:

$$\frac{\partial p}{\partial x} = \mu \frac{\partial^2 u_x}{\partial y^2} \tag{3}$$

According to Equation 3, greater convexity in the velocity profile along the *y*-axis increases the pressure difference along the *x*-direction. When coating blade velocity was set to 10 mm $s^{-1}$, the rGO-chitosan solution was found to have a relatively linear velocity profile between 0 and 45 μm in height (along the *y*-axis) compared to other coating blade velocity conditions. We may therefore deduce that the pressure difference near the blade would be the smallest for when coating speed is 10 mm $s^{-1}$, which showed the most linear velocity change.

**Supplementary Videos description**

**Video S1.** Top-view in situ optical microscopy of the rGO-chitosan solution solidification process after the blade has traversed across the substrate with a coating speed of 3 mm $s^{-1}$

**Video S2**. Top-view in situ optical microscopy of the rGO-chitosan solution solidification process after the blade has traversed across the substrate with a coating speed of 9 mm $s^{-1}$

**Video S3.** Top-view in situ optical microscopy of the rGO-chitosan solution solidification process after the blade has traversed across the substrate with a coating speed of 60 mm $s^{-1}$

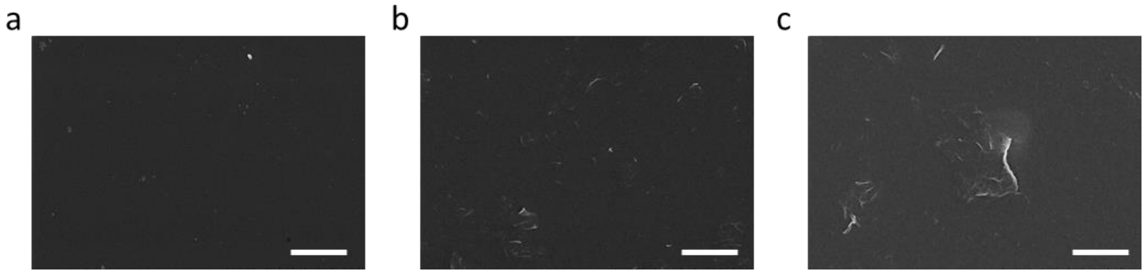


**Figure S1.** SEM images of the rGO-based nanocomposites at different coating temperatures **a,** 70ºC **b,** 80ºC, and **c,** 90ºC. The rGO-chitosan solution consisted of 0.2 mg $ml^{-1}$ rGO and 10 mg $ml^{-1}$ chitosan solution and were coated at 0.5 mm $s^{-1}$. Scale bars are 2 μm.

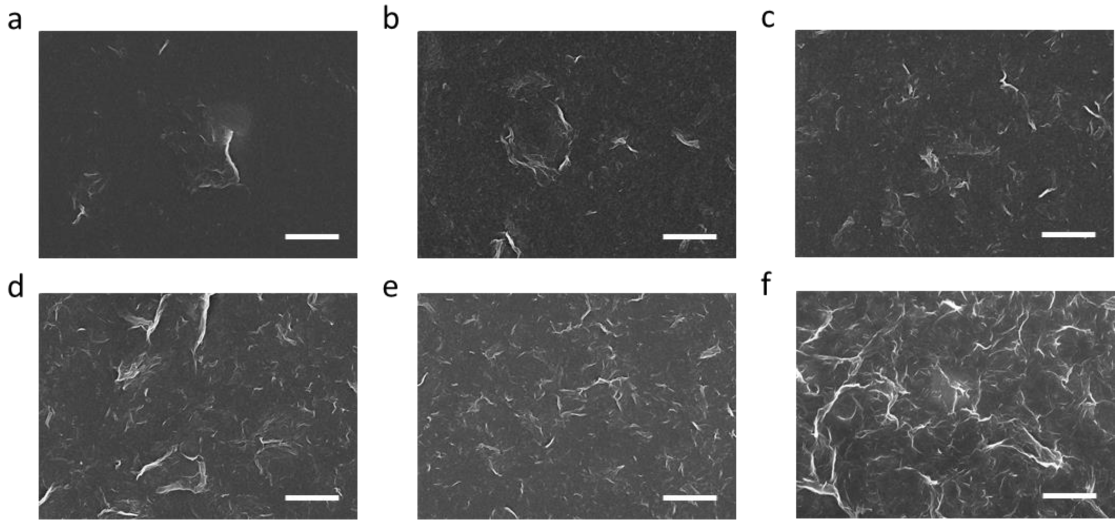


**Figure S2.** SEM images of rGO-based nanocomposites produced with different rGO to chitosan ratios, where chitosan concentrations are fixed at 10 mg $mL^{-1}$, and rGO concentrations are **a,** 0.1 mg $mL^{-1}$ **b,** 0.5 mg $mL^{-1}$ **c,** 1 mg $mL^{-1}$ **d,** 2 mg $mL^{-1}$ **e,** 3 mg $mL^{-1}$, and **f,** 8 mg $mL^{-1}$. Scale bars are 2 μm.

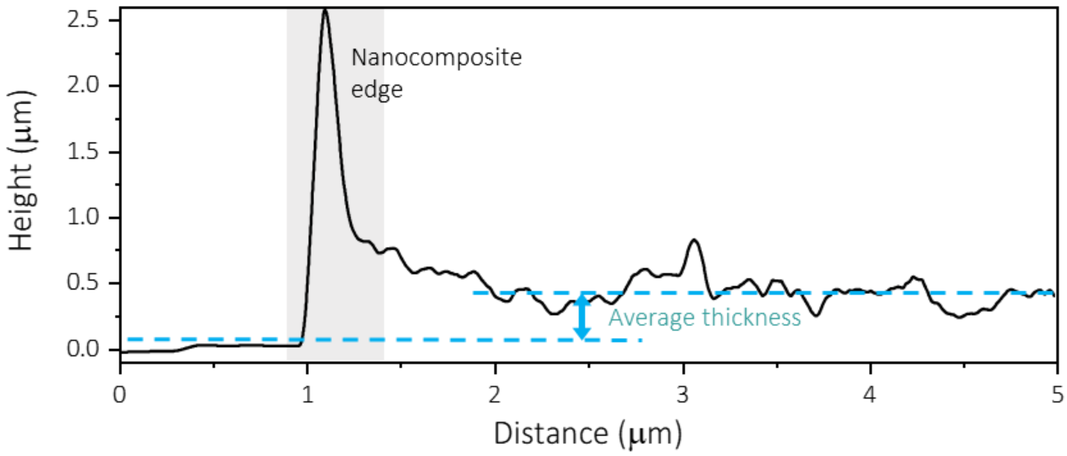


**Figure S3.** Thickness calculation method represented with the rGO-based nanocomposite coated at 0.3 mm $s^{-1}$. The difference between average height values before and after the nanocomposite edge (gray) identifiable through a distinct peak is calculated as the average thickness.

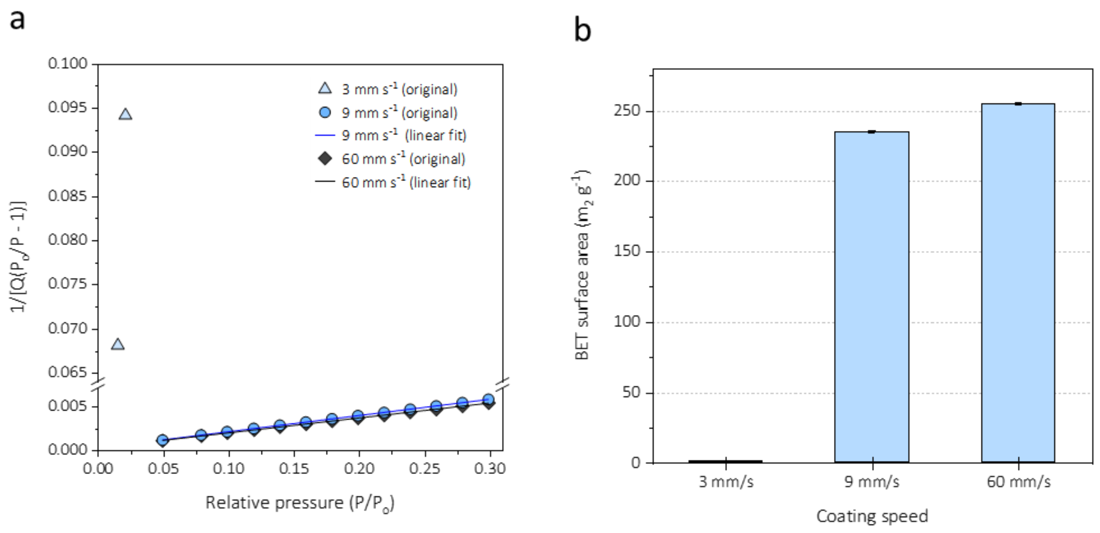


**Figure S4. a,** BET surface area plot of the rGO-based nanocomposite at different coating speeds (3, 9, and 60 mm $s^{-1}$). High correlation coefficient value indicates the high reliability of BET surface area. Correlation coefficient of 9 mm $s^{-1}$ and 60 mm $s^{-1}$ is 1.00, and 3 mm $s^{-1}$ does not fit. **b,** BET surface area values of the rGO-based nanocomposite at different coating speeds (3, 9, and 60 mm $s^{-1}$). BET surface area of 3 mm $s^{-1}$ is 1.0045 $m^2$ $g^{-1}$, BET surface area of 9 mm $s^{-1}$ is 234.7843 $m^2$ $g^{-1}$, and BET surface area of 70 mm $s^{-1}$ is 254.3706 $m^2$ $g^{-1}$.

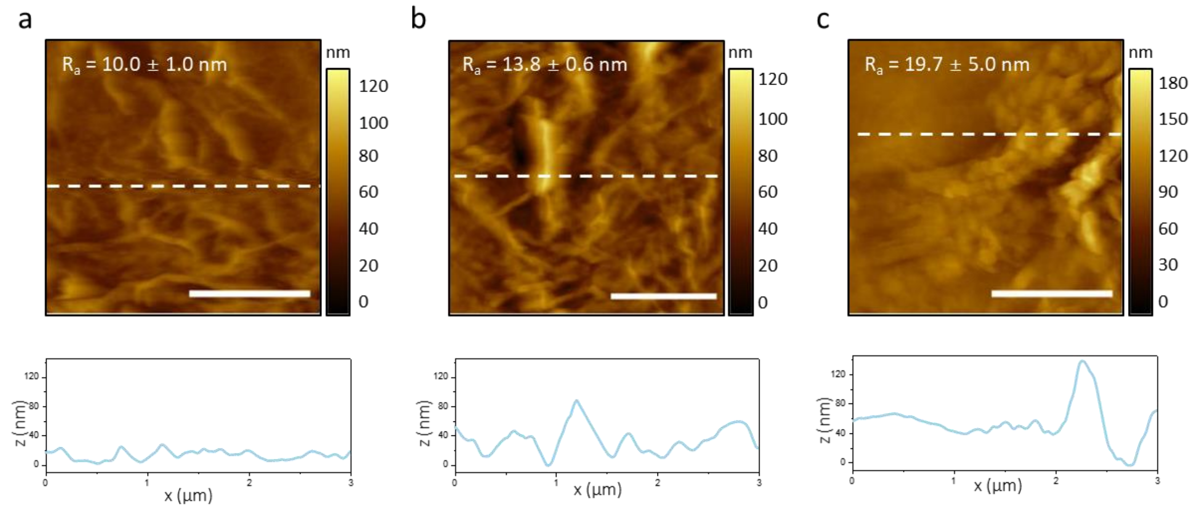


**Figure S5.** AFM images with average roughness and their corresponding surface profiles (white dashed lines) of the rGO-based nanocomposites produced at different coating speeds. **a,** 3 mm $s^{-1}$ shows small thickness and low degree of wrinkling, **b,** 9 mm $s^{-1}$ shows small thickness and high degree of wrinkling, and **d,** 60 mm $s^{-1}$ shows high thickness and unevenly distributed wrinkles and aggregates. Scale bars are 1 μm. Average roughness and standard deviation were determined by 4 different samples.

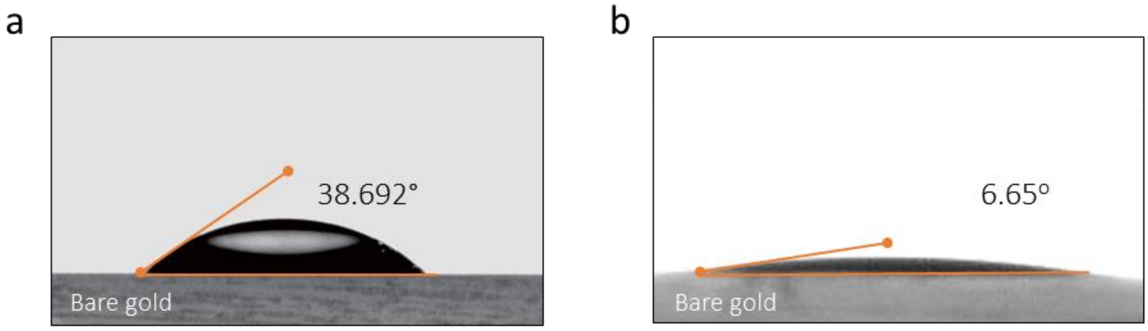


**Figure S6.** Contact angle analysis by optical images of **a,** a droplet of rGO suspension in DI-water as control and **b,** rGO-chitosan solution droplet on bare gold electrodes. Drop volume of each solution was 5 ul. Contact angle of rGO-chitosan solution on bare gold surface showed a 5.82-fold decrease compared to control, indicating a more favorable interaction between the electrode surface and the rGO-chitosan solution.

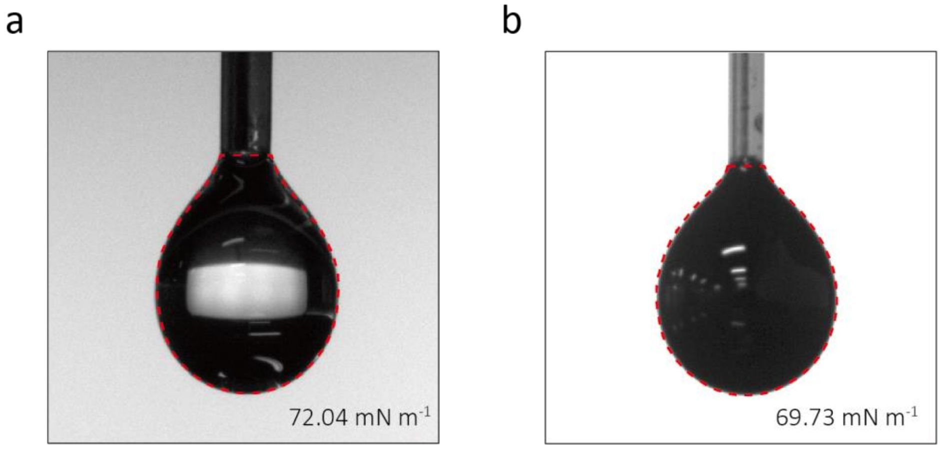


**Figure S7.** Surface tension measurement of the **a,** droplet of rGO suspension in DI-water as control (volume=11.55 μL) and **b,** rGO-chitosan solution (volume=11.29 μL) by pendant drop analysis. Red dashed line represents the fitting line for measurement. The drop of the rGO-chitosan solution shows a surface tension value of 69.73 mN $m^{-1}$ while that of control shows a value of 72.04 mM $m^{-1}$.

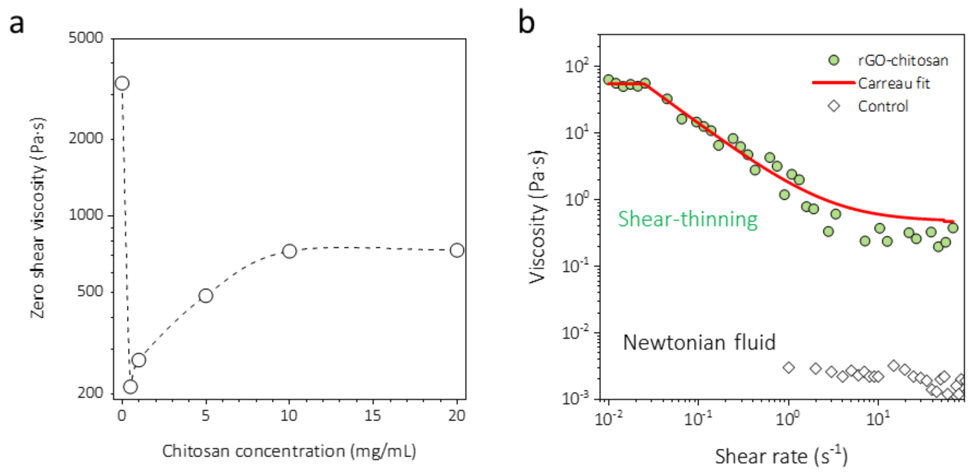


**Figure S8.** Rheological properties of the rGO-chitosan solution **a,** Zero-shear viscosity of rGO-chitosan solutions with changing chitosan concentration from 0 to 20 mg $mL^{-1}$, and rGO concentration fixed at 5 mg $mL^{-1}$. Zero-shear viscosity value was saturated at 10 mg $mL^{-1}$ of chitosan, which indicates that the viscosity initially increased by the physical tangling and molecular interactions between chitosan and rGO molecules. **b,** Measurement of viscosity against shear rate showing the shear-thinning behavior of rGO-chitosan solution composed of 5 mg $mL^{-1}$ rGO and 10 mg $mL^{-1}$ chitosan. A solid red line is fitted with the Carreau model indicating a non-Newtonian model. Meanwhile, an aqueous solution used as a control shows a Newtonian model.

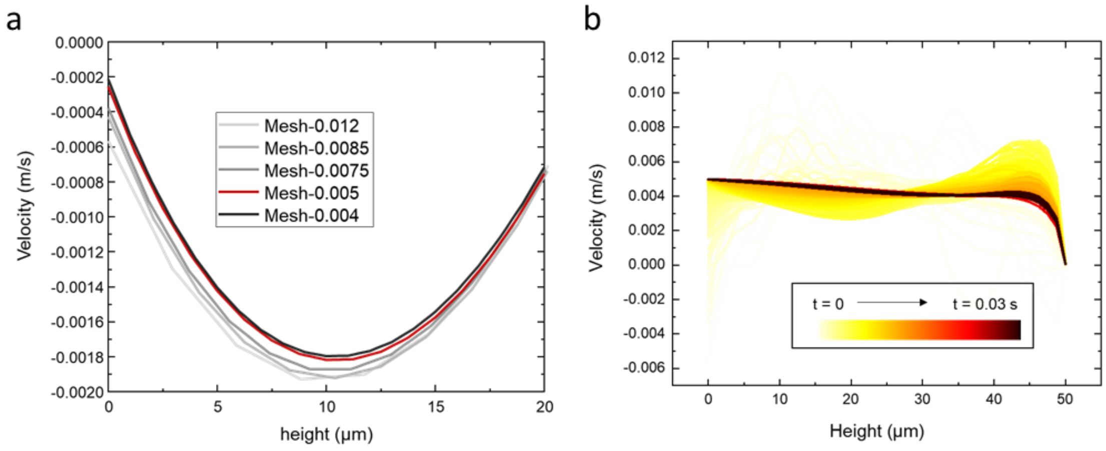


**Figure S9.** Convergence of an LSM simulation. **a,** The convergence of rGO-chitosan solution velocity at Inside by mesh size when the velocity of the blade (U) is 20 mm $s^{-1}$. Each result is labeled as “Mesh-k”, where the value of k represents maximum element size coefficient based along the +*y* direction at Inside. Higher k value refers a coarser mesh near the entrance of the blade. Mesh-0.005 was chosen for the simulation**. b,** The convergence of rGO-chitosan solution velocity at Inside in a function of time (t) when U is 20 mm $s^{-1}$. Darker colors indicate higher t. The timespan for simulation was set to 0.03 s.

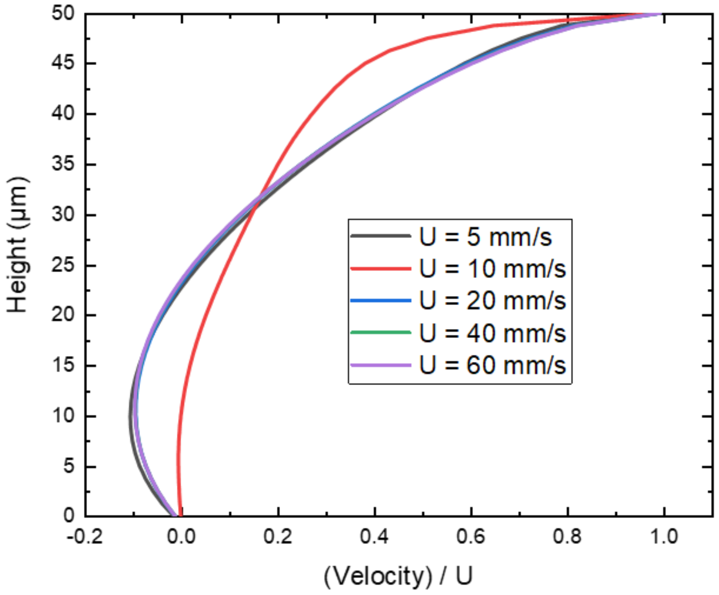


**Figure S10.** The normalized rGO-chitosan solution velocity profile along the +y-direction at Inside. Coating blade velocity of 10 mm $s^{-1}$ exhibits the minimum decrease in velocity profile while others show greater velocity changes.

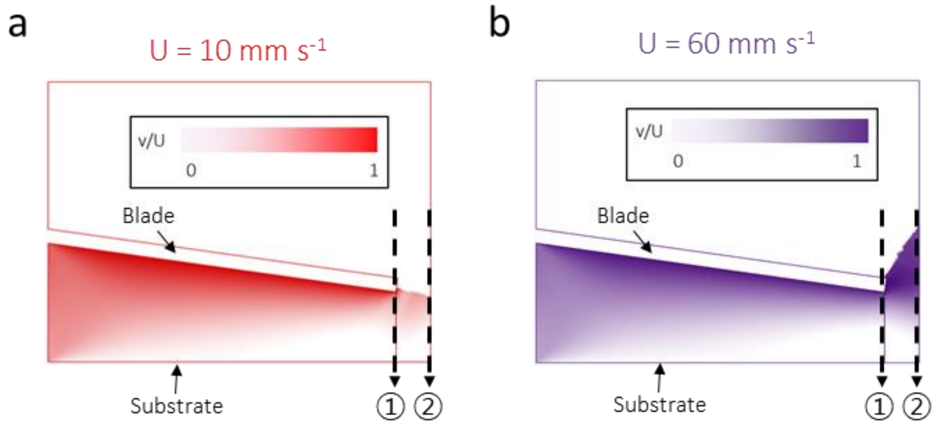


**Figure S11.** Prediction of fluid velocity during solution shearing process and fluid thickness of Outside at **a,** the velocity of 10 mm $s^{-1}$ and **b,** the velocity of 60 mm $s^{-1}$. The edge indicated by '1' refers to the starting point of the coating (Inside), and the edge indicated by '2' refers to the finishing point of the coating (Outside), where the blade has moved 25 µm in the x-axis direction.

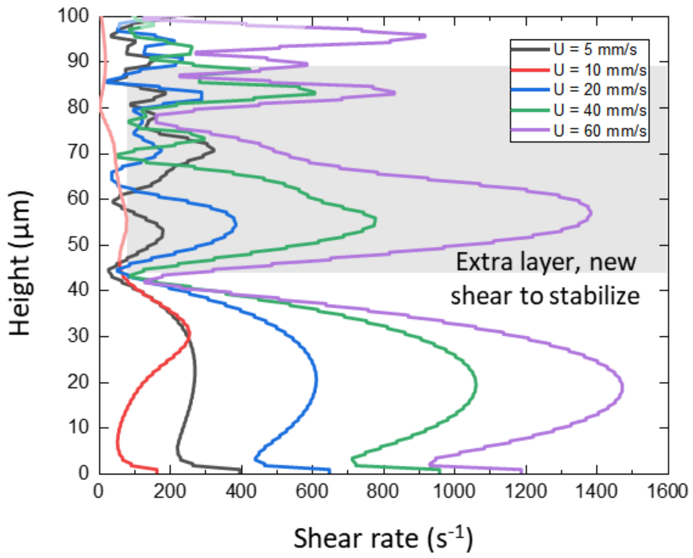


**Figure S12.** Shear rate profiles of the rGO-chitosan solution along the +*y*-direction at Outside while the velocity of blade increases from 5 mm $s^{-1}$ to 60 mm $s^{-1}$. In the case of 5, 20, 40, and 60 mm $s^{-1}$, the thickness of the fluid after blade movement turns out to be thicker than the gap between blade and the substrate, generating extra fluid layers. This results in a longer stabilization time compared to that of 10 mm $s^{-1}$.

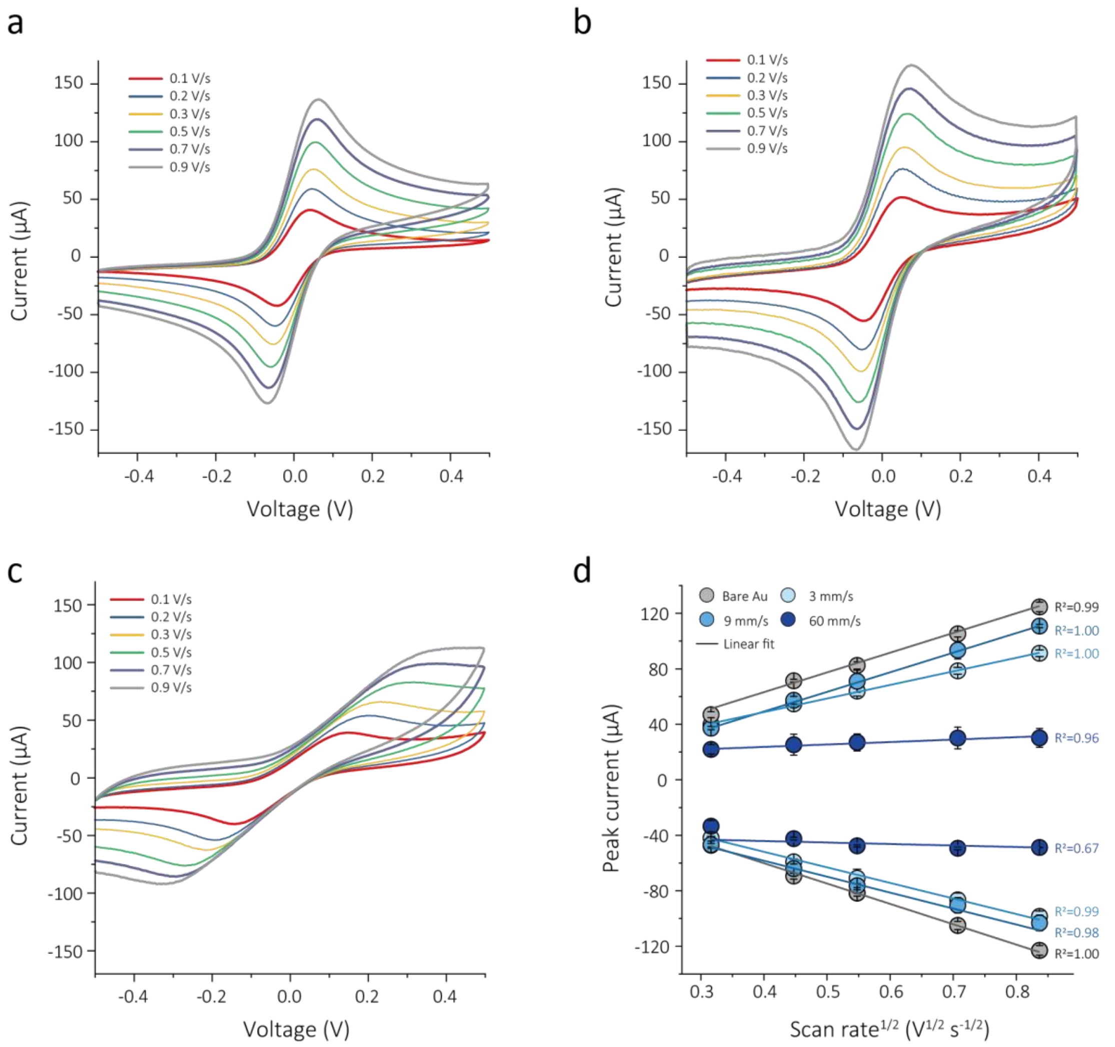


**Figure S13.** Cyclic voltammetry profiles by different scan rates for the rGO-based nanocomposites produced at **a,** 3 mm $s^{-1}$ **b,** 9 mm $s^{-1}$, and **c,** 60 mm $s^{-1}$. **d,** Oxidation and reduction current densities of bare gold electrode and rGO-based nanocomposites produced at three coating speeds against the square root of scan rate. (n=4, independent electrode, error bars=standard deviation). The linearity reflects the dependence of this composite's electrochemical sensing mechanism on electron diffusion. The nanocomposite at 9 mm $s^{-1}$ scan rate shows a steepness closely matching that of bare gold, meaning the nanocomposite allows a similar level of free electron diffusion. The decreasing steepness for the nanocomposite made at 3 mm $s^{-1}$ indicates a decrease in this freedom due to the reduced number of wrinkles. A weak proportionality at 60 mm $s^{-1}$ points to quasi-reversible electrochemical reaction.

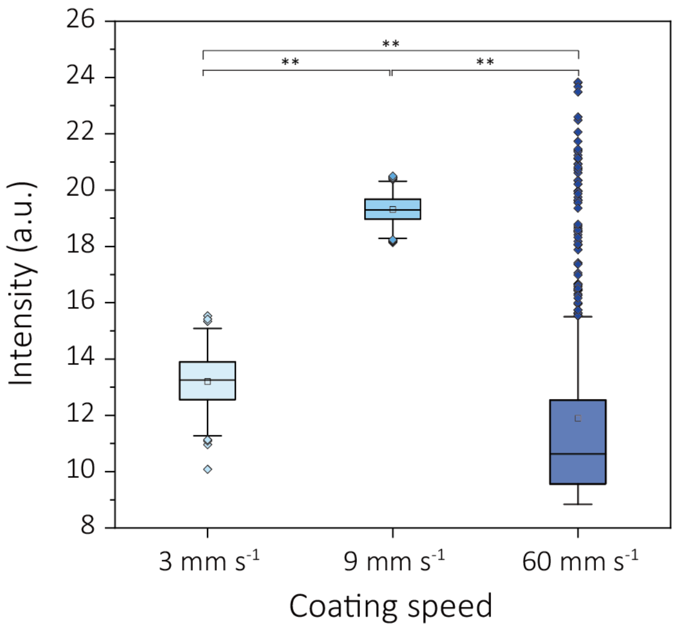


**Figure S14.** FITC fluorescence intensities extracted using image J from the confocal images of the rGO-based nanocomposites produced at three coating speeds (3, 9, and 60 mm $s^{-1}$). Each nanocomposite was covalently immobilized with FITC-labeled anti-IgG. Average intensity at 9 mm $s^{-1}$ shows a CV of 2.43%, while 3 mm $s^{-1}$ and 60 mm $s^{-1}$ show CVs of 7.04% and 27.80%, respectively. Statistical significance was tested (**$P < 0.01$; one-way ANOVA and tukey post hoc test).

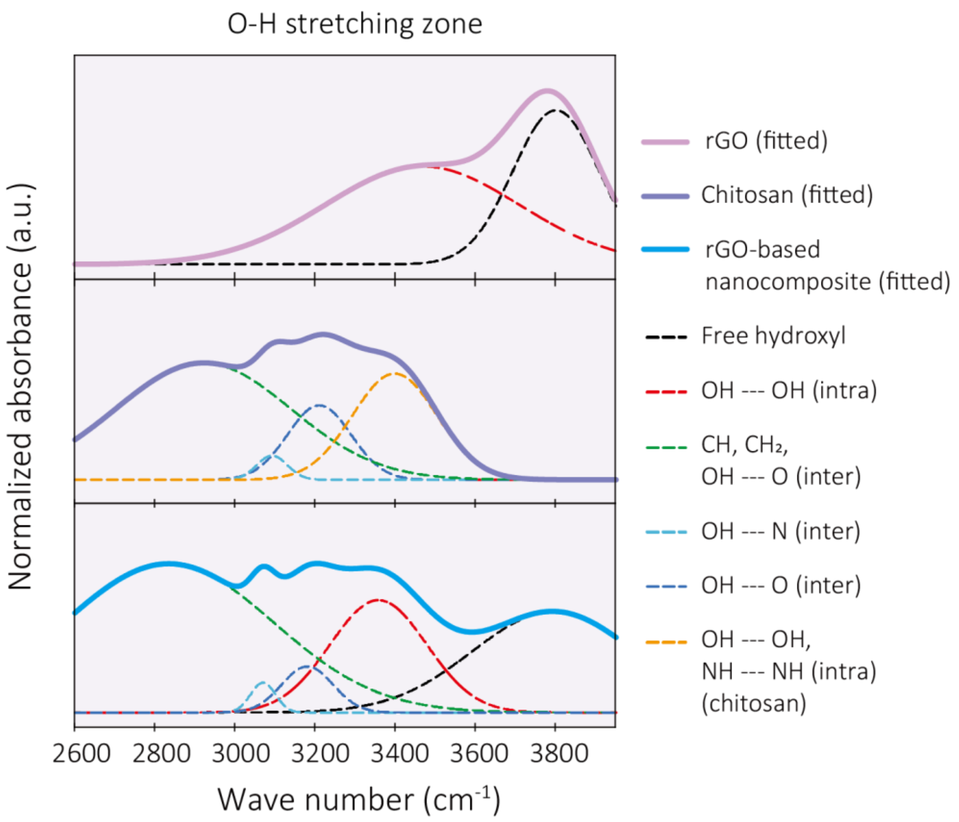


**Figure S15.** Deconvolution of O-H stretching peaks found between 2600 and 4100 $cm^{-1}$ in the normalized absorbance FT-IR spectra of rGO, chitosan, and the optimized rGO-based nanocomposite (9 mm $s^{-1}$). Each nonlinear fitted line shows $R^2$ values of 0.994, 0.990, and 0.997, respectively.

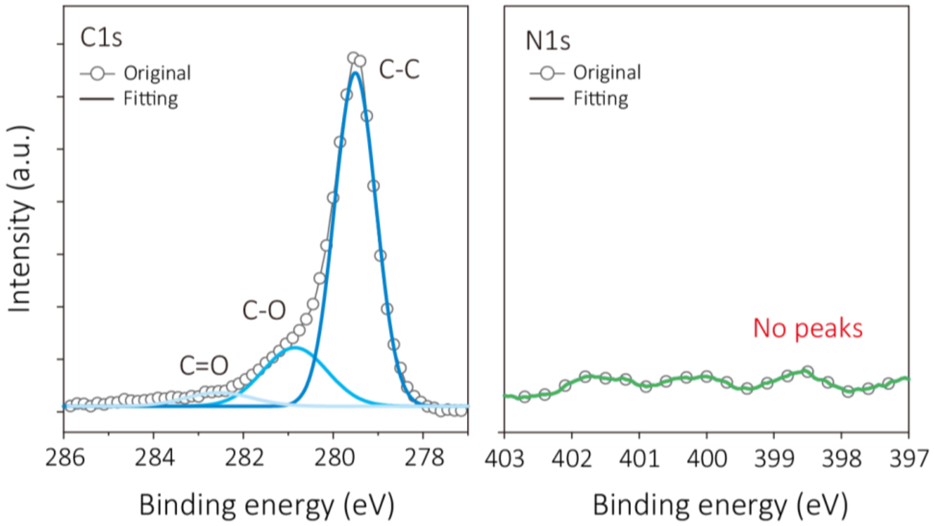


**Figure S16.** XPS spectra and its deconvoluted peaks for core C1s (left) and N1s (right) from the rGO film.

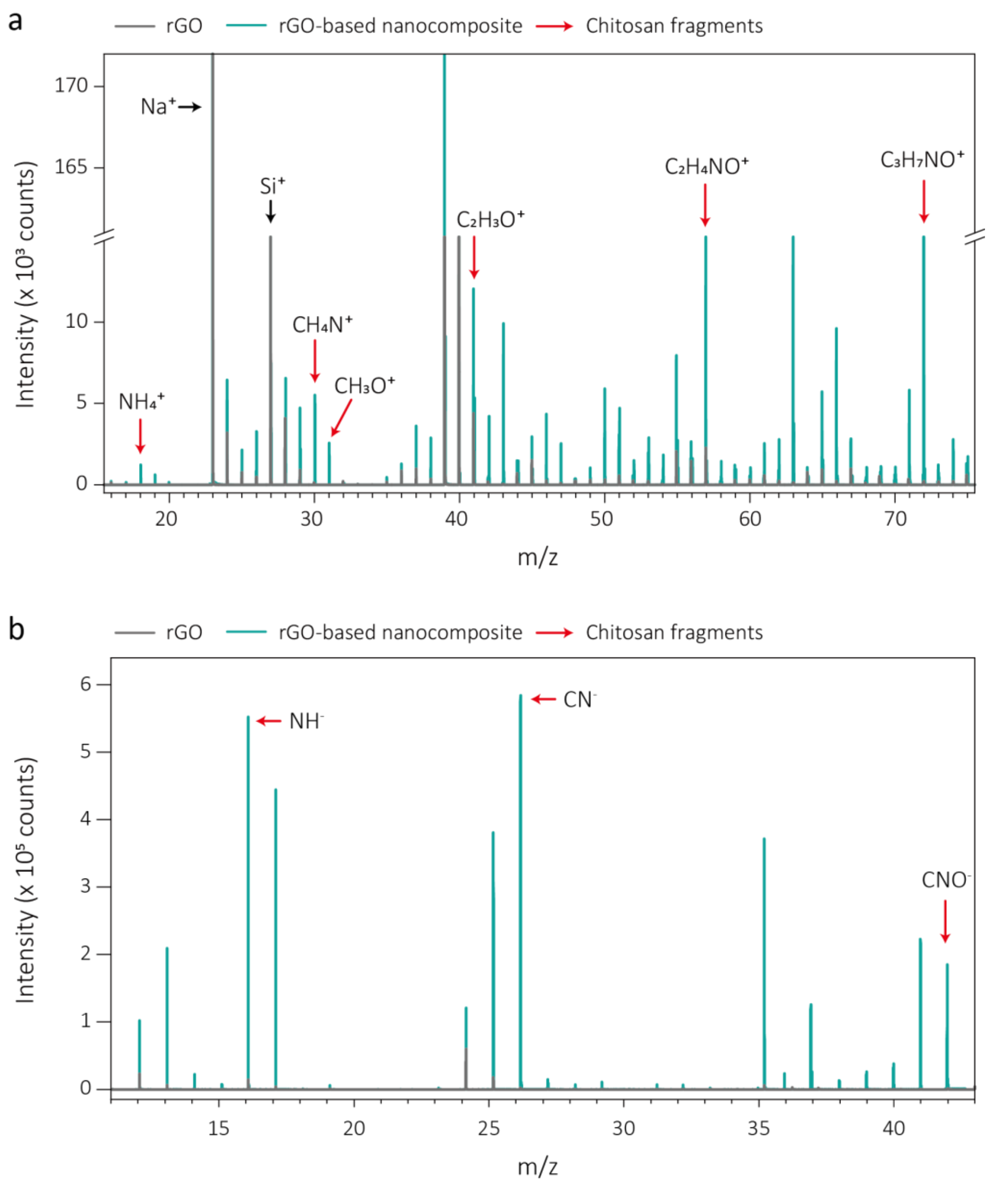


**Figure S17.** TOF-SIMS spectra of the rGO film and the optimized rGO-based nanocomposite (9 mm $s^{-1}$). *y*-axis indicates the detected number of secondary ions and *x*-axis indicates the mass-to-charge (m/z) ratio. **a,** In positive TOF-SIMS spectra, unlike the rGO film, the rGO-based nanocomposite reveals various fragments derived from chitosan fragments, such as $NH_4^+$, $CH_4N^+$, $CH_3O^+$. $C_2H_3O^+$, $C_2H_4O^+$, and $C_3H_7NO^+$. **b,** In negative TOF-SIMS spectra, chitosan fragments are evident in the rGO-based nanocomposite from the peaks corresponding to $NH^-$, $CN^-$, and $CNO^-$, which are not present in the rGO films. These results reflect the successful incorporation of chitosan with rGO molecules.

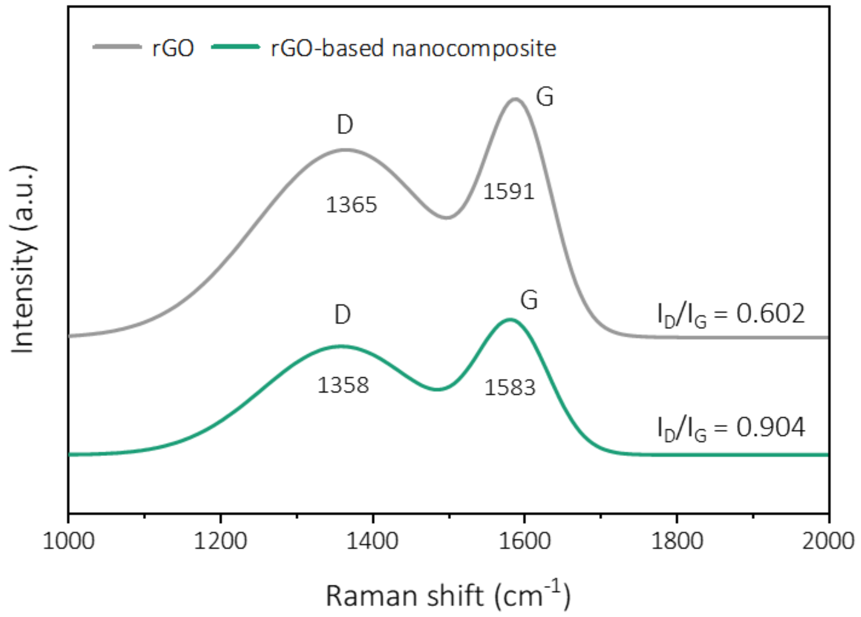


**Figure S18.** Raman spectra of the rGO-film and the optimized rGO-based nanocomposite (9 mm s$^{-1}$) both showing distinct D ($E_{2g}$ vibrational mode of sp$^2$ carbon) and G ($A_{1g}$ symmetric vibrational mode) bands. The increase in D/G intensity ratio upon the addition of chitosan indicates increased number of defects, reflecting their successful incorporation between rGO.

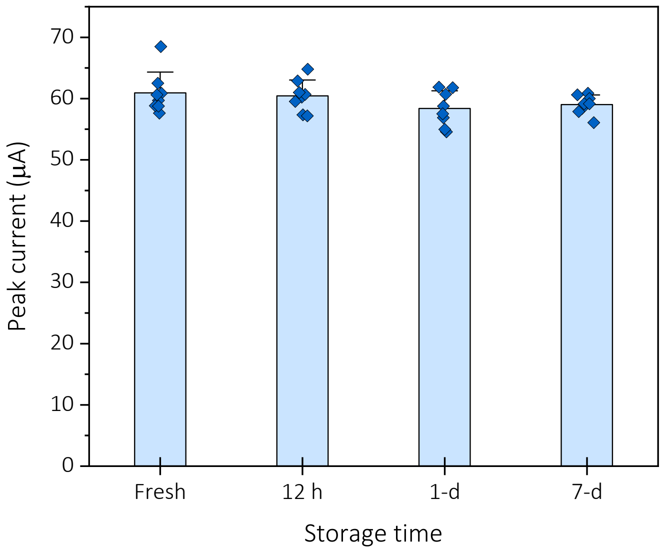


**Figure S19.** Shelf-life of the rGO-based nanocomposite chip stored in 10 mM PBS at 25 °C. (n=8, independent electrode, error bars=standard deviation)

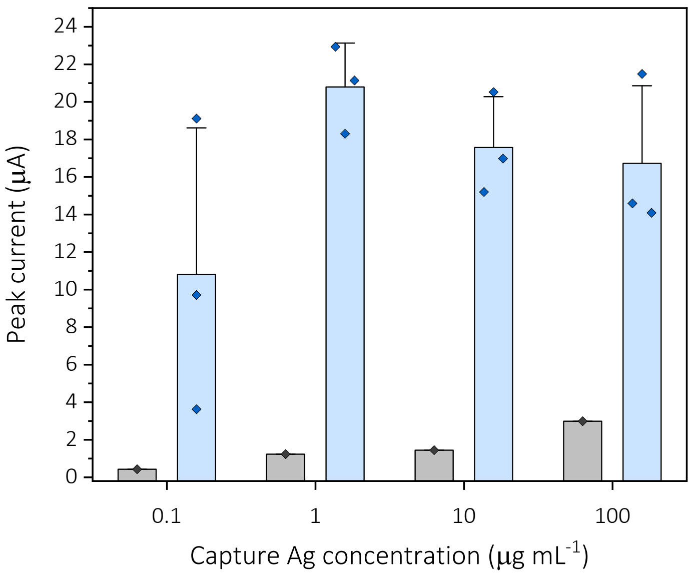


**Figure S20.** Optimization of capture protein concentration for antibody detection using the rGO-based nanocomposite chip (n=3, independent electrode, error bars=standard deviation) with PDIA6 as capture protein, 10 ng $mL^{-1}$ of anti-PDIA6, and 50 μg $mL^{-1}$ of HRP-labeled detection antibody. Gray bars indicate signals from negative control, incubated in BSA.

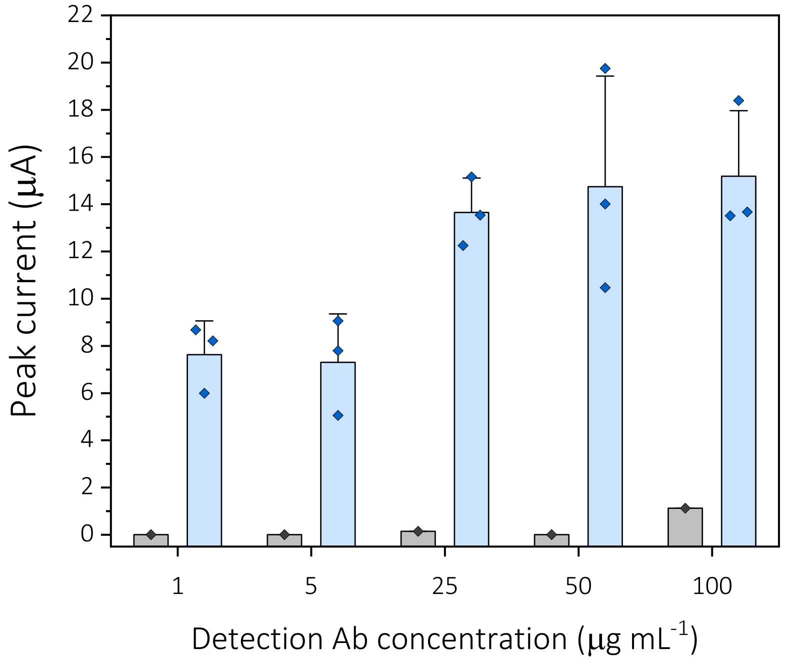


**Figure S21.** Optimization of HRP-labeled detection antibody concentration for antibody detection using the rGO-based nanocomposite chip (n=3, independent electrode, error bars=standard deviation). With HRP-labeled detection antibody, 1 μg $mL^{-1}$ of capture protein (PDIA6), and 10 ng $mL^{-1}$ of anti-PDIA6 antibody. Gray bars indicate negative control, incubated in BSA.

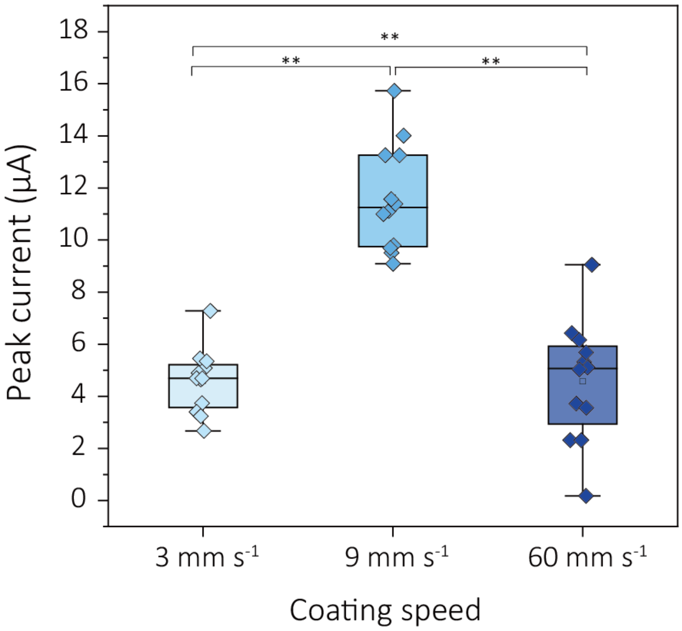


**Figure S22.** Electrochemical signal response of the rGO-based nanocomposites coated at different coating speeds (3, 9, and 60 mm $s^{-1}$). Electrodes immobilized with capture proteins (PDIA6), specifically bound with 10 ng $mL^{-1}$ of anti-PDIA6 and HRP-labeled detection antibodies underwent TMB precipitation, followed by cyclic voltammetry measurements in 10 mM PBS. (n=12, independent electrode, error bars=standard deviation). Statistical significance was tested (**P < 0.01; one-way ANOVA and tukey post hoc test).

| Experimental set | anti-PDIA6 | anti-PERK | anti-GRP78 |
|---|---|---|---|
| 1 | 0.1 ng $mL^{-1}$ | | 100 ng $mL^{-1}$ |
| 2 | 1 ng $mL^{-1}$ | | 50 ng $mL^{-1}$ |
| 3 | 10 ng $mL^{-1}$ | 10 ng $mL^{-1}$ | 10 ng $mL^{-1}$ |
| 4 | 50 ng $mL^{-1}$ | | 1 ng $mL^{-1}$ |
| 5 | 100 ng $mL^{-1}$ | | 0.1 ng $mL^{-1}$ |
| 6 | 100 ng $mL^{-1}$ | 0.1 ng $mL^{-1}$ | |
| 7 | 50 ng $mL^{-1}$ | 1 ng $mL^{-1}$ | |
| 8 | 10 ng $mL^{-1}$ | 10 ng $mL^{-1}$ | 10 ng $mL^{-1}$ |
| 9 | 1 ng $mL^{-1}$ | 50 ng $mL^{-1}$ | |
| 10 | 0.1 ng $mL^{-1}$ | 100 ng $mL^{-1}$ | |
| 11 | | 100 ng $mL^{-1}$ | 0.1 ng $mL^{-1}$ |
| 12 | | 50 ng $mL^{-1}$ | 1 ng $mL^{-1}$ |
| 13 | 10 ng $mL^{-1}$ | 10 ng $mL^{-1}$ | 10 ng $mL^{-1}$ |
| 14 | | 1 ng $mL^{-1}$ | 50 ng $mL^{-1}$ |
| 15 | | 0.1 ng $mL^{-1}$ | 100 ng $mL^{-1}$ |

**Table S1. Experimental sets for multiplexed sensing.** Each target sample is prepared by spiking a PBS solution with each of the three antibodies in the concentrations specified above. The sensor results for experimental sets 1-5, 6-10, and 11-15 correspond to Figure 5f, g, and h, respectively.
